# Homeostatic Dysregulation of Systemic CD8^+^ T Cell Compartment in Lung Cancer Patients

**DOI:** 10.1101/2023.12.27.573474

**Authors:** Sung-Woo Lee, Ju Sik Yun, Young Ju Kim, Hee-Ok Kim, Hyun-Ju Cho, Cheol-Kyu Park, In-Jae Oh, Jae-Ho Cho

## Abstract

Cancer adapts various resistance mechanisms to counteract CD8^+^ T cell attacks. While this suppression of antigen-specific CD8^+^ T cells is common within the tumor microenvironment, little is known about how tumors affect CD8^+^ T cells systemically. Here we show a new link between tumor-associated homeostatic dysregulation and uncontrolled differentiation of peripheral blood CD8^+^ T cells. These CD8^+^ T cells exhibited progressive alterations indicative of diminished quiescence, increased spontaneous activation, and more-differentiated proliferation-incompetent effector cells. This phenomenon was not limited to tumor-reactive cells but broadly applicable to non-specific cells, correlating with poor clinical responses to immune checkpoint inhibitor therapy. These findings provide a new mechanism by which cancer impairs CD8^+^ T cells by dysregulating the homeostasis of systemic CD8^+^ T cell populations.

**One-Sentence Summary:** Cancer-associated homeostatic dysregulation accelerates uncontrolled differentiation of systemic CD8^+^ T cells.

## Main Text

It has long been recognized that the immune system can detect, monitor, and eliminate cancer, and today, numerous studies have made this even more evident. For example, the correlation between an increase in tumor-infiltrating lymphocytes (especially CD8^+^ T cells) and responsiveness to immunotherapy is known to indicate a cancer-immune interaction(*1-4*). Additionally, various markers suggestive of an active anti-tumor immune response within tumor tissues (e.g., PD-L1 expression, tumor mutation burden, interferon (IFN)-γ production) have been reported with their associations with clinical responses to immunotherapy(*3-13*). However, paradoxically, the presence of anti-tumor immunity highlights the fact that cancer can evolve immune resistance as a protective adaptation against immune attack. Indeed, among various evasion mechanisms, IFN-γ produced by tumor-reactive CD8^+^ T cells is thought to inhibit anti-tumor activity by inducing PD-L1 expression within tumor tissues(*14-16*). Immune checkpoint inhibitors (ICIs) targeting PD-1/PD-L1 inhibit such cancer-induced adaptive resistance mechanisms and, therefore, have revolutionized cancer treatment(*4-7, 17-19*). However, they still have significant limitations in that they are only effective in ∼20% of cancer patients(*20-22*). Furthermore, the fact that ∼60% of non-responders to anti-PD-(L)1 therapy belong to a group with high tumor PD-L1 expression(*19*) suggests that there are much broader and more complex immune evasion mechanisms beyond immune checkpoint–mediated suppression in the tumor microenvironment.

Tumor-specific CD8^+^ T cells have shown to be heterogeneous in their activation and differentiation status, with a correlation with clinical responses to ICI therapy(*23-32*). Such heterogeneity is considered to reflect various contexts of anti-tumor immune responses, but the precise nature of these contexts (both within and outside of tumor microenvironment) and their possible relevance to cancer-associated immune resistance mechanisms are still poorly understood. Despite growing knowledge of the mechanisms of cancer-associated resistance to tumor-reactive CD8^+^ T cells and the clinical applications of this understanding(*33, 34*), the overall survival of cancer patients has not significantly improved(*35-37*). In this study, we sought to address the question of whether cancer can affect the immunosurveillance function of CD8^+^ T cells at a systemic level and explore its underlying mechanisms. We conducted an extensive retrospective analysis of peripheral blood CD8^+^ T cells from 349 lung cancer patients, and found an aberrantly gross alteration in the homeostatic regulation of the systemic CD8^+^ T cell compartment, which causes uncontrolled differentiation from proliferation-competent to proliferation-incompetent effector subsets, leading to poor clinical responses to ICI therapy. Therefore, we uncovered a new evasion mechanism by which cancer avoids attack by potentially tumor-reactive CD8^+^ T cells.

## RESULTS

### Progressive alterations in peripheral blood CD8^+^ Tem of lung cancer patients

To investigate whether tumors induce alterations in the composition of peripheral blood CD8^+^ T cells, we retrospectively analyzed CD8^+^ T cells in the peripheral blood mononuclear cells (PBMCs) of 53 healthy donors and 349 lung cancer patients (294 non-small cell lung cancer (NSCLC) and 55 small cell lung cancer (SCLC) patients; table S1). Based on the expression of CCR7 and CD45RA (Fig. 1A), CD8^+^ T cells were classified into naïve (Tn; CCR7^+^CD45RA^+^), central memory (Tcm; CCR7^+^CD45RA^−^), effector memory (Tem; CCR7^−^CD45RA^−^), and effector memory re-expressing CD45RA (Temra; CCR7^−^CD45RA^+^). Interestingly, patients exhibited a higher frequency of Tem than healthy individuals (Fig. 1, B and C), and this observation remained consistent in age-matched comparisons (fig. S1, A and B). Such an increase was frequently seen in healthy older individuals but was notably significant even among the younger patients (Fig. 1D).

**Fig. 1.**
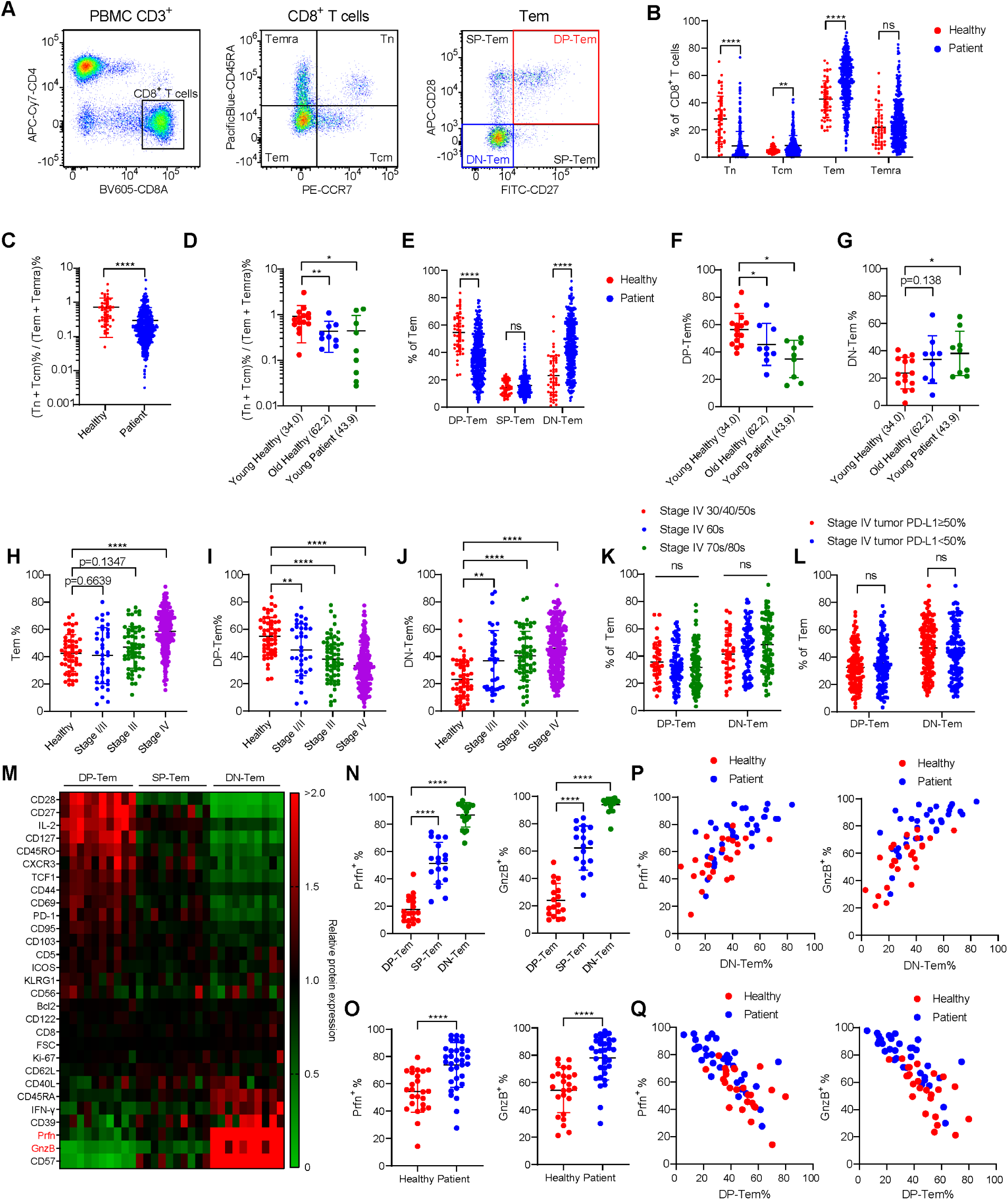
Lung cancer patients show altered peripheral blood CD8^+^ Tem subsets. (**A**) Gating strategies for CD8^+^ T cell subsets. (**B**) CD8^+^ T cell subset frequencies in healthy individuals (n=53) and patients (n=349). (**C**–**D**) Ratio between less-differentiated and more-differentiated subsets. Numbers in brackets represent the mean age of the group. (**E**) CD8^+^ Tem subset frequencies. (**F**–**G**) (F) DP-Tem and (G) DN-Tem frequencies in Tem population. (**H**–**J**) (H) Tem, (I) DP-Tem and (J) DN-Tem frequencies in patients at different stages of lung cancer. (**K**–**L**) DP-Tem and DN-Tem frequencies in stage IV patients grouped by (K) age or (L) tumor PD-L1 expression. (**M**) Relative protein expressions of various molecules in Tem subsets (n=10). Relative expressions were calculated by dividing MFIs with that of total Tem. Expressions were color-coded from green (low) to red (high). (**N**–**O**) Prfn^+^ and gnzB^+^ frequencies. (**P**–**Q**) Correlation between prfn^+^ and gnzB^+^ frequencies and (P) DN-Tem or (Q) DP-Tem frequencies. All bar graphs represent mean ± SD, ∗∗∗∗p<0.0001, ∗∗∗p<0.001, ∗∗p<0.01, ∗p<0.05.

Moreover, when Tem were further subdivided by CD27 and CD28 expression (Fig. 1A, right), patients showed a lower frequency of less-differentiated CD27^+^CD28^+^ double-positive (DP)-Tem and higher frequency of more-differentiated CD27^−^CD28^−^ double-negative (DN)-Tem compared to healthy individuals (Fig. 1E). This was consistently observed in age-matched comparisons (Fig. 1, F and G, and fig. S1C) and, to a lesser extent, in comparisons of DP- and DN-Temra (fig. S1, D to F).

Next, we investigated whether the elevated frequency of more-differentiated peripheral blood CD8^+^ Tem among cancer patients was associated with tumor progression. There was a stage-dependent gradual rise in the frequency of Tem (but not Tn and Tcm), accompanied by a decrease in DP-Tem frequency and an increase in DN-Tem frequency in NSCLC patients (Fig. 1, H to J, and fig. S1, G and H). A similar trend was observed among Temra (fig. S1, I to K). Notably, the decreased DP-Tem frequency and increased DN-Tem frequency in stage IV NSCLC patients was also seen in extensive disease (ED) stage SCLC patients (fig. S1L), but was irrelevant to age, tumor PD-L1 expression, sex, histologic classification, or EGFR mutation (Fig. 1, K and L, and fig. S1, M to O).

We further explored whether the above changes in CD8^+^ Tem subpopulations were associated with functional changes. Differential expression of various proteins was observed during DP-to-DN transition with a notable increase in the expression of cytotoxic molecules, perforin (prfn) and granzyme B (gnzB) (Fig. 1, M and N). Importantly, the high expression of prfn/gnzB in Tem was strongly correlated with the increased frequency of DN-Tem; in fact, Tem of patients exhibited higher prfn/gnzB expression than those of healthy individuals (Fig. 1O). Furthermore, this phenomenon showed a strong positive correlation with DN-Tem frequency and a negative correlation with DP-Tem frequency (Fig. 1, P and Q). These findings imply cancer-associated potential alterations in the phenotypic and functional composition of systemic CD8^+^ T cell populations, especially Tem, during tumor progression.

### Effect of ICI therapy on DP to DN transition in CD8^+^ Tem

We investigated whether the observed alterations in peripheral blood CD8^+^ Tem subpopulations were indeed associated with anti-tumor immune responses. For this, we retrospectively analyzed the differentiation states and compositional changes of CD8^+^ T cells using PBMCs isolated from the blood of 32 NSCLC patients collected before (pre) and after (post) anti-PD-L1 therapy (Fig. 2A and table S2). There was a subtle increase in the frequency of Tem post-therapy (Fig. 2B and fig. S2A). However, interestingly, anti-PD-L1 therapy led to substantial alterations in the frequency of DP- and DN-Tem (Fig. 2, C to E); ∼71.9% of the patients exhibited reduced DP-Tem frequency, while ∼68.8% exhibited increased DN-Tem frequency post-therapy (Fig. 2E). Moreover, the frequency of prfn^+^/gnzB^+^ Tem also increased post-therapy (Fig. 2F), consistent with the positive correlation between DN-Tem frequency and prfn^+^/gnzB^+^ frequency (Fig. 1P). Furthermore, patients who exhibited significantly high fold change (FC) increases in prfn^+^/gnzB^+^ frequency between pre- and post-therapy (FC^hi^, FC≥1.1; Fig. 2G) demonstrated better clinical responses than those who exhibited low or no increases in FC (FC^lo^, FC<1.1; Fig. 2H).

**Fig. 2.**
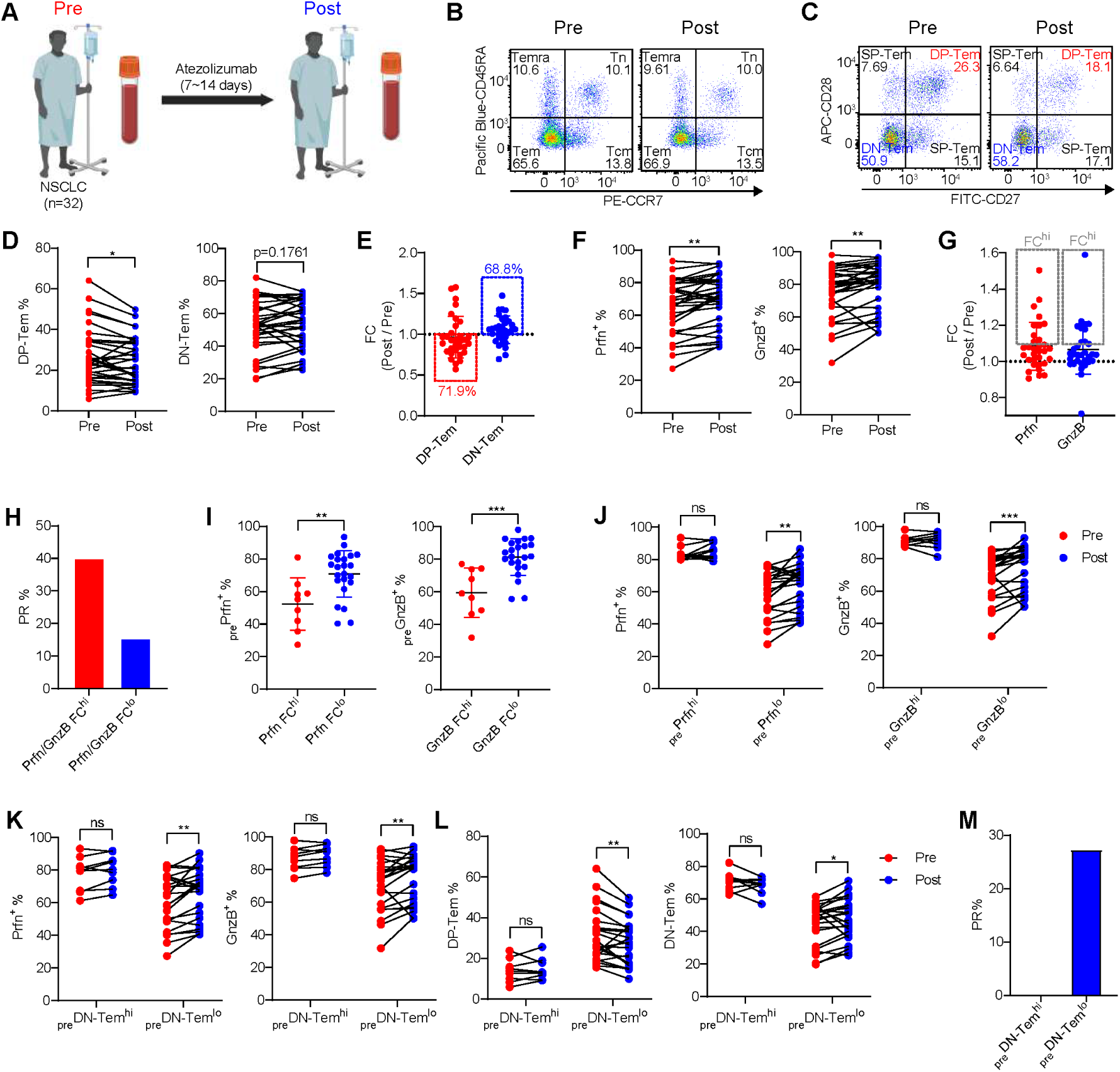
ICI therapy induces altered CD8^+^ Tem subsets. (**A**) Illustration of the blood collection process. (**B**–**C**) Representative flow cytometry data comparing (B) CD8^+^ T cell subset and (C) Tem subset frequencies. (**D**–**E**) (D) Changes in DP-Tem and DN-Tem frequencies after ICI therapy and (E) their fold changes (FC) (n=32). (**F**–**G**) (F) Changes in prfn^+^ and gnzB^+^ frequencies after ICI therapy and (G) their FC. Grey box indicates patients with high FC (FC^hi^). (**H**) Proportion of patients with PR to ICI therapy in Prfn/GnzB FC^hi^ (Prfn FC^hi^ and GnzB FC^hi^) and Prfn/GnzB FC^lo^ (Prfn FC^lo^ and/or GnzB FC^lo^) groups. (**I**) _pre_Prfn^+^ and _pre_GnzB^+^ frequencies. (**J**) Changes in prfn^+^ (or gnzB^+^) frequencies after ICI therapy. (**K**–**L**) Changes in (K) prfn^+^ and gnzB^+^ frequencies and (L) DP-Tem and DN-Tem frequencies after ICI therapy. (**M**) Proportion of patients with PR to ICI therapy. All bar graphs represent mean ± SD, ∗∗∗p<0.001, ∗∗p<0.01, ∗p<0.05.

Given the higher therapeutic responses of FC^hi^ patients compared with FC^lo^ patients, we sought to determine an explanation for this difference. Remarkably, we observed significant differences in the baseline prfn^+^/gnzB^+^ (_pre_Prfn^+^/_pre_GnzB^+^) frequency between FC^hi^ and FC^lo^ patients (Fig. 2I). Patients with low (lo) _pre_Prfn^+^/_pre_GnzB^+^ frequency (_pre_Prfn^lo^/_pre_GnzB^lo^) had significant increases in the post-therapy prfn^+^/gnzB^+^ frequency, whereas no such increase was observed in patients with high (hi) _pre_Prfn^+^/_pre_GnzB^+^ frequency (_pre_Prfn^hi^/_pre_GnzB^hi^) (Fig. 2J and fig. S2B). Moreover, consistent with the positive correlation (Fig. 1P), similar increases were observed in patients with relatively lower (but not higher) pre-therapy DN-Tem frequency (_pre_DN-Tem^lo^ vs _pre_DN-Tem^hi^; Fig. 2K), and this phenomenon was accompanied by decreased post-therapy DP-Tem frequency and increased DN-Tem frequency (Fig. 2L). Furthermore, _pre_DN-Tem^lo^ patients showed better therapeutic responses than _pre_DN-Tem^hi^ patients (Fig. 2M and fig. S2C). A similar phenomenon was observed in _pre_DP-Tem^hi^ but not _pre_DP-Tem^lo^ patients (fig. S2, D to G).

Based on the above findings, the decreased _pre_DP-Tem and increased _pre_DN-Tem frequency in the patients’ blood was interpreted as a negative factor that diminishes anti-tumor CD8^+^ T cell responses to ICI therapy. In support of this, significantly reduced proliferative responses were observed in DN-Tem relative to DP-Tem upon TCR stimulation *in vitro*, and likewise, increases of the post-therapy Ki-67^+^ cells were far smaller in DN-Tem than DP-Tem (fig. S2, H and I). Collectively, these findings suggest that the cancer-associated DP-to-DN-Tem transition in NSCLC patients may reflect sustained, recurrent anti- or perhaps pro-tumor immunity, potentially impacting therapeutic responses to ICI therapy.

### Effect of clonal expansion on the increased CD8^+^ DN-Tem

We investigated the mechanism underlying the increased frequency of DN-Tem in NSCLC patients. For this, CD8^+^ T cells were isolated from PBMCs of four healthy individuals and eight NSCLC patients (table S3). This was followed by scRNA-seq, scTCR-seq and CITE-seq for analyzing gene expression profile, TCR repertoire, and CD45RA protein expression, respectively (Fig. 3A). We classified 13 clusters (C0–C12) using an unsupervised clustering method (Fig. 3B). Among these, C0 and C1 clusters were designated as *C0.Tn* and *C1.Tcm*, based on high expression of Tn signature genes (*CCR7, LEF1, SELL, TCF7, ACTN1*) and Tcm signature genes (*GPR183, GATA3, CRIP2*), respectively, along with CD45RA protein expression (Fig. 3, C and D, and fig. S3, A and B). C2 cluster was defined as *C2.DP-Tem* based on its relatively high expression of early-effector cell signature genes (*CD27, CD28, GZMK*) and low CD45RA protein expression (Fig. 3, C and D, and fig. S3, A and B). Unlike the C2 cluster, C3–C9 clusters exhibited high expression of late-effector cell signature genes (*GZMA, GZMB, GZMH, PRF1, CX3CR1, PLEK, NKG7, KLRD1, B3GAT1, KLRG1*; Fig. 3, C and D). These clusters showed somewhat heterogeneous CD45RA expression, with levels between those of Tem and Temra (fig. S3, A and B), but for simplicity, they were collectively designated as *C3–9.DN-Tem*. Among the C3–C9 clusters, C3 cluster was observed in all individuals analyzed, while C4–C9 clusters exhibited highly limited *TCRα* and *TCRβ* sequences and were exclusively observed in specific individuals only (Fig. 3, C and E); therefore, we designated them as “common” *C3.DN-Tem* (*C3.DN-Tem-CO*) and “individual-specific” *C4–9.DN-Tem* (*C4–9.DN-Tem-IS*), respectively. Additionally, C10, C11, and C12 clusters showed high expression of long non-coding RNA genes (*NEAT1, MALAT1*), S100 genes (*S100A8, S100A9, S100A10*), and MAIT cell signature genes (*KLRB1, CEBPD, LTK, CXCR6, CCR6*), which were designated as *C10.lncRNA high*, *C11.S100 high*, and *C12.MAIT*, respectively (Fig. 3, B and C, and fig. S3C).

**Fig. 3.**
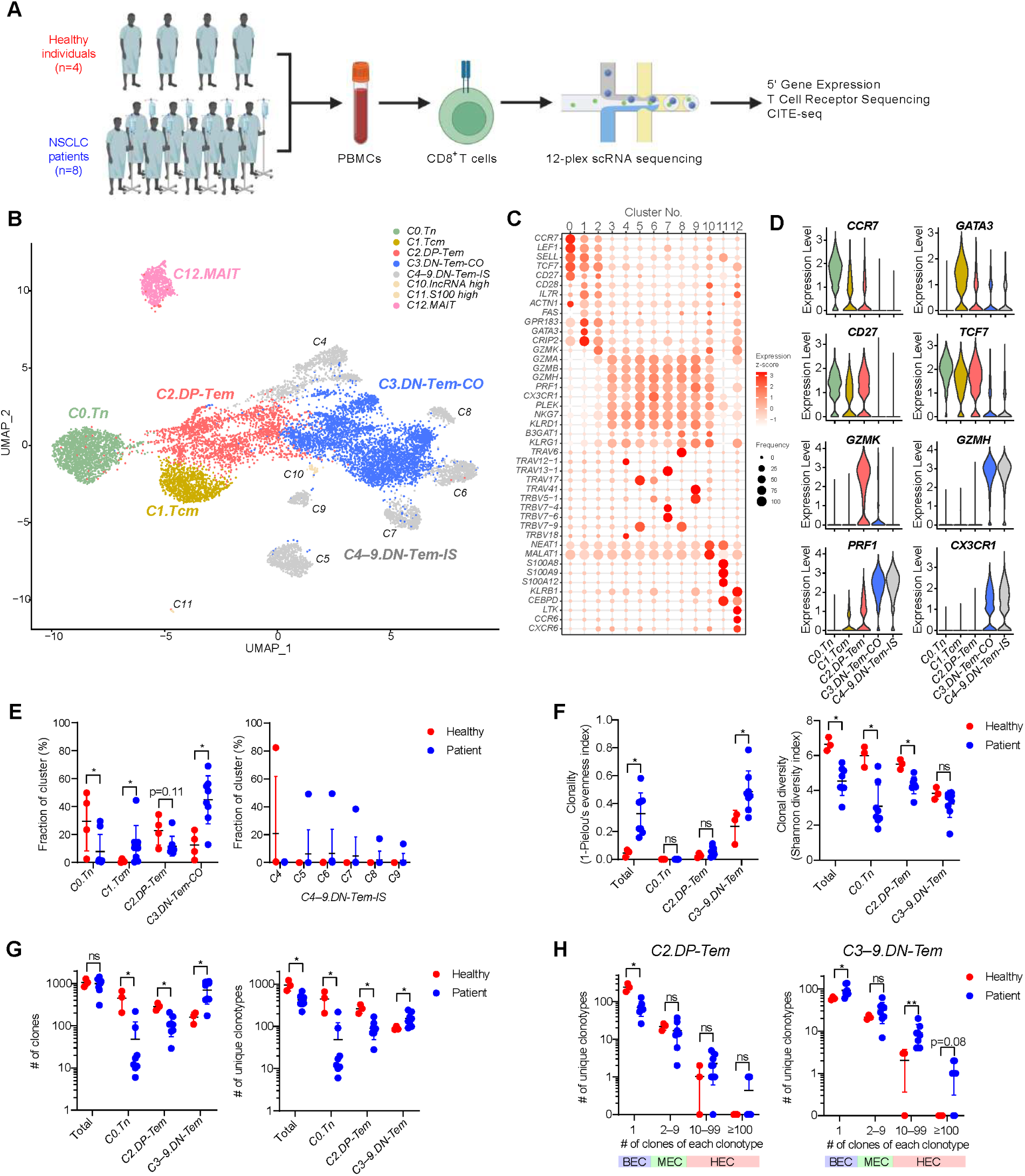
Increased CD8^+^ DN-Tem in NSCLC patients rely on both clonal expansion-dependent and -independent mechanisms. **(A)** Experimental scheme for scRNA-seq. **(B)** UMAP of scRNA-seq. Clusters were color-coded according to their labels. **(C)** Expressions of signature genes in each cluster. Color and size represent relative average expression (z-score) and frequency, respectively. **(D)** Expression of key signature genes in *C0–9* clusters. **(E)** Frequencies of *C0–9* clusters. (**F**–**G**) (F) Clonality, clonal diversity, (G) numbers of clones and unique clonotypes in indicated clusters. (**H**) The numbers of unique clonotypes with BEC, MEC, and HEC. All bar graphs represent mean ± SD, ∗∗p<0.01, ∗p<0.05.

With reference to the C0–C9 clusters, NSCLC patients had lower proportions of *C0.Tn* and *C2.DP-Tem* and higher proportions of *C3.DN-Tem-CO* and *C4–9.DN-Tem-IS* than healthy individuals (Fig. 3E), consistent with the previous flow cytometry data (Fig. 1, B and E). Among *C4–9.DN-Tem-IS*, *C4* cluster showed exceptionally rare occurrence in only one (H2) of the four healthy individuals (Fig. 3E, right); therefore, H2 was excluded from further analysis. Next, we performed TCR repertoire analysis to ascertain whether the elevated proportion of *DN-Tem* observed in NSCLC patients involved oligoclonal expansion. As expected, the clonality of the mixture of clusters (C0–C9; representing total peripheral blood CD8^+^ T cells) significantly increased in patients compared with healthy individuals, while their diversity substantially decreased (see “Total”; Fig. 3F). The reduced diversity was particularly prominent in the *C0.Tn* and *C2.DP-Tem*, but notably was absent in the *C3–9.DN-Tem* (Fig. 3F, right), despite their increased clonality (Fig. 3F, left).

Moreover, the numbers of both clones (i.e., counting all individual clonotypes, including identical clonotypes) and unique clonotypes (i.e., counting each unique individual clonotype) in *C3–9.DN-Tem* were significantly increased in patients relative to healthy individuals (Fig. 3G). Particularly, the increase in the number of unique clonotypes was observed not only in heavily expanded clones (HEC: ≥10 clones) but also notably in barely expanded clones (BEC: 1 clone), albeit no differences in moderately expanded clones (MEC: 2–9 clones) (Fig. 3H, right). This phenomenon in *C3–9.DN-Tem* was particularly evident in *C3.DN-Tem-CO* but not *C4–9.DN-Tem-IS* (fig. S3, D to F). In contrast, the numbers of both clones and unique clonotypes in *C0.Tn* and *C2.DP-Tem* was significantly reduced in patients relative to healthy individuals (Fig. 3G), and the reduction in the number of unique clonotypes was observed only in BEC (Fig. 3H, left, and fig. S3G). These results indicate that peripheral blood CD8^+^ DN-Tem frequency in NSCLC patients increases without causing a loss in their diversity despite elevated clonality, indicating a possible role of a clonal expansion-independent mechanism.

### Diversification of CD8^+^ DN-Tem into *GZMK^+^.DN-Tem versus GZMK^−^.DN-Tem*

We investigated how the above unexpected clonal expansion-independent increase of DN-Tem arises. For this, we further characterized the *C3.DN-Tem-CO* based on pseudotime and compared any potential differences between patients and healthy individuals. Interestingly, a marked difference was apparent especially at relatively earlier pseudotime points for *C3.DN-Tem-CO*, as evidenced by a significantly higher fraction of cells in patients than in healthy individuals (Fig. 4A; grey box). These cells observed in the *C3.DN-Tem-CO* expressed *GZMH* at high levels and interestingly also expressed *GZMK* albeit in lower amounts (Fig. 4, B and C). Therefore, *C3–9.DN-Tem* were further characterized into two clusters: *GZMK^+^.C3–9.DN-Tem* and *GZMK^−^.C3–9.DN-Tem* (Fig. 4D). *GZMK^+^.C3–9.DN-Tem* was distributed between *C2.DP-Tem* and *GZMK^−^.C3–9.DN-Tem* (Fig. 4D, yellow in the UMAP) and expressed both early (*GZMK, CD27, TCF7*) and late effector genes (*KLRD1, GZMH, CX3CR1*) (Fig. 4E). Notably, *GZMK^+^.C3–9.DN-Tem* was nearly absent in healthy individuals but significantly elevated in patients (Fig. 4, F and G). More importantly, the increased number of unique clonotypes observed in *C3–9.DN-Tem* (Fig. 3G) was primarily attributable to *GZMK^+^.C3–9.DN-Tem* rather than *GZMK^−^.C3–9.DN-Tem* (Fig. 4, H and I).

**Fig. 4.**
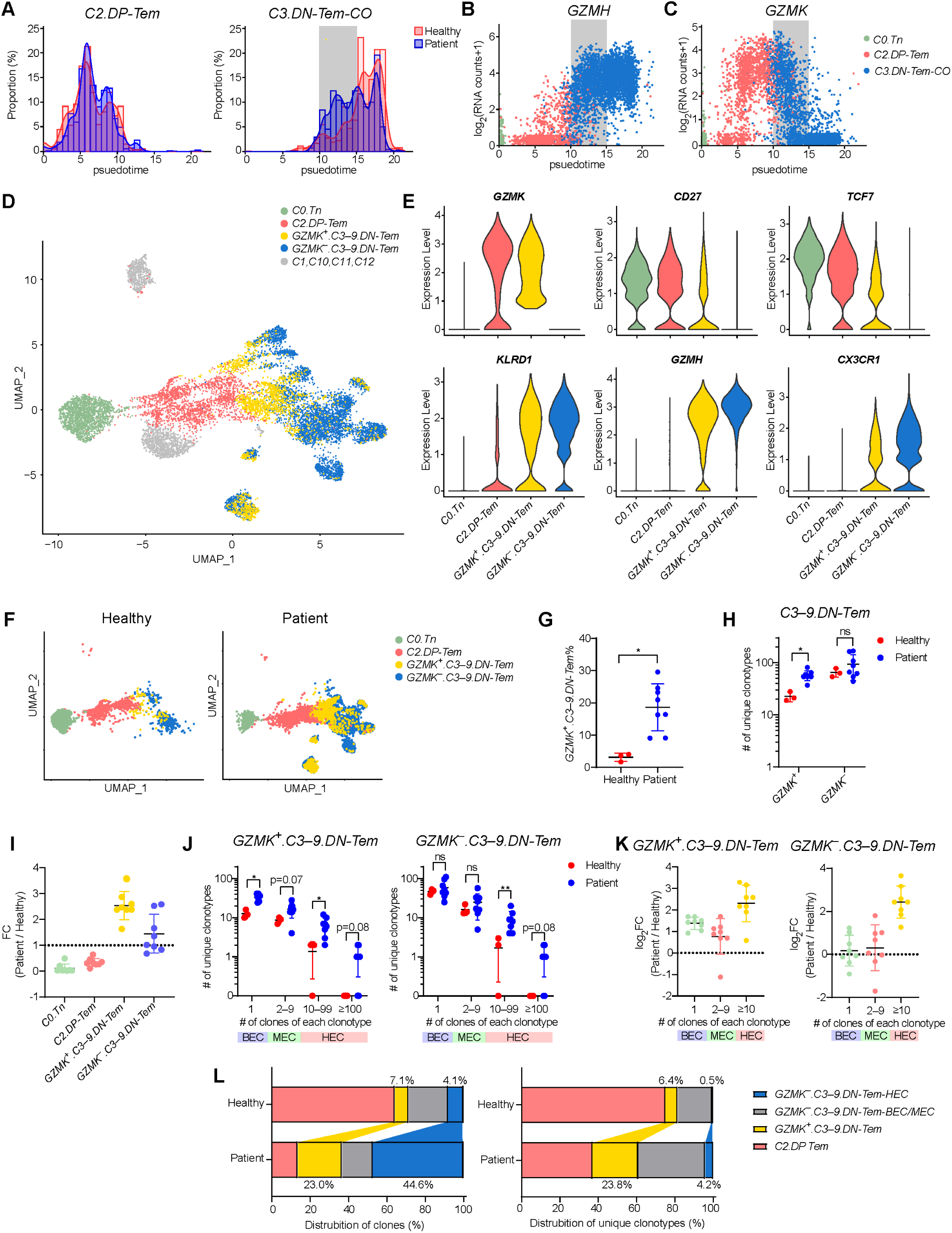
Increased CD8^+^ DN-Tem in NSCLC patients are associated with clonally diverse *GZMK^+^.DN-Tem* generation. (**A**) Histogram for pseudotime of *C2.DP-Tem* and *C3.DN-Tem-CO*. The grey box indicates the pseudotime where patients and healthy individuals were different. (**B**–**C**) (B) GZMH and (C) GZMK expression by pseudotime. (**D**) UMAP highlighting *GZMK^+^.C3–9.DN-Tem*. (**E**) Expressions of early- and late-effector signature genes. (**F**) Separate UMAP for healthy individuals and patients. (**G**) *GZMK^+^.C3–9.DN-Tem* frequency in CD8^+^ T cells. (**H**–**I**) (H) Number of unique clonotypes (I) and their FC. (**J**–**K**) (J) Number of unique clonotypes with BEC, MEC, and HEC (K) and their FC. (**L**) Distribution of clones and unique clonotypes in Tem (*C2–9*) clusters. All bar graphs represent mean ± SD, ∗∗p<0.01, ∗p<0.05.

In our further analysis, the increased number of unique clonotypes observed in the *GZMK^+^.C3–9.DN-Tem* was evident across all clonotypes with HEC, MEC, and even BEC (Fig. 4, J and K, left). It should be noted, however, that *GZMK^−^.C3–9.DN-Tem*, despite the lack of difference shown in Fig. 4H, also showed a significant increase especially in clonotypes only with HEC (Fig. 4, J and K, right). Based on these findings, we again further categorized *C3–9.DN-Tem* into three subclusters—*GZMK^−^.C3–9.DN-Tem-HEC, GZMK^−^.C3–9.DN-Tem-BEC/MEC, and GZMK^+^.C3–9.DN-Tem*—to examine the relative distribution of either clones or unique clonotypes within each subcluster (Fig. 4L). *GZMK^+^.C3–9.DN-Tem* had the most significant contribution to both the clone and unique clonotype distribution in patients compared to healthy individuals, whereas *GZMK^−^.C3–9.DN-Tem-HEC* only contributed to clone distribution but had minimal effects on the unique clonotype distribution (Fig. 4L). All these observations support the notion that peripheral blood CD8^+^ Tem in NSCLC patients undergo a continuous DP-to-DN-Tem transition, which is important for shaping clonal diversity within DN-Tem, including an increase in *GZMK^+^.DN-Tem* that is independent of clonal expansion.

### Transcriptional signatures associated with reduced T cell quiescence

The increased frequency of clonally diverse *GZMK^+^.DN-Tem* in NSCLC patients raises the question of whether this increase is solely due to tumor antigen-specific responses. In fact, when compared to the 75,302 known virus-specific CDR3 sequences in VDJdb(*38*), a fraction of *GZMK^+^.DN-Tem* (particularly *GZMK^+^.C3.DN-Tem-CO*) contained clonotypes restricted to known viral epitopes (fig. S4A), suggesting that not all *GZMK^+^.DN-Tem* are tumor specific. We therefore further investigated the mechanism of how tumor non-specific *GZMK^+^.DN-Tem* could be increased in NSCLC patients. Interestingly, significant differences in gene expression profiles were observed in *C2.DP-Tem*, *C0.Tn*, *C3.DN-Tem*, and even *C12.MAIT*, as evidenced by clearly distinguishable positions in the UMAP (Fig. 5A). This suggests that tumor-specific transcriptomic changes occur broadly and systemically across various cell types in peripheral blood.

**Fig. 5.**
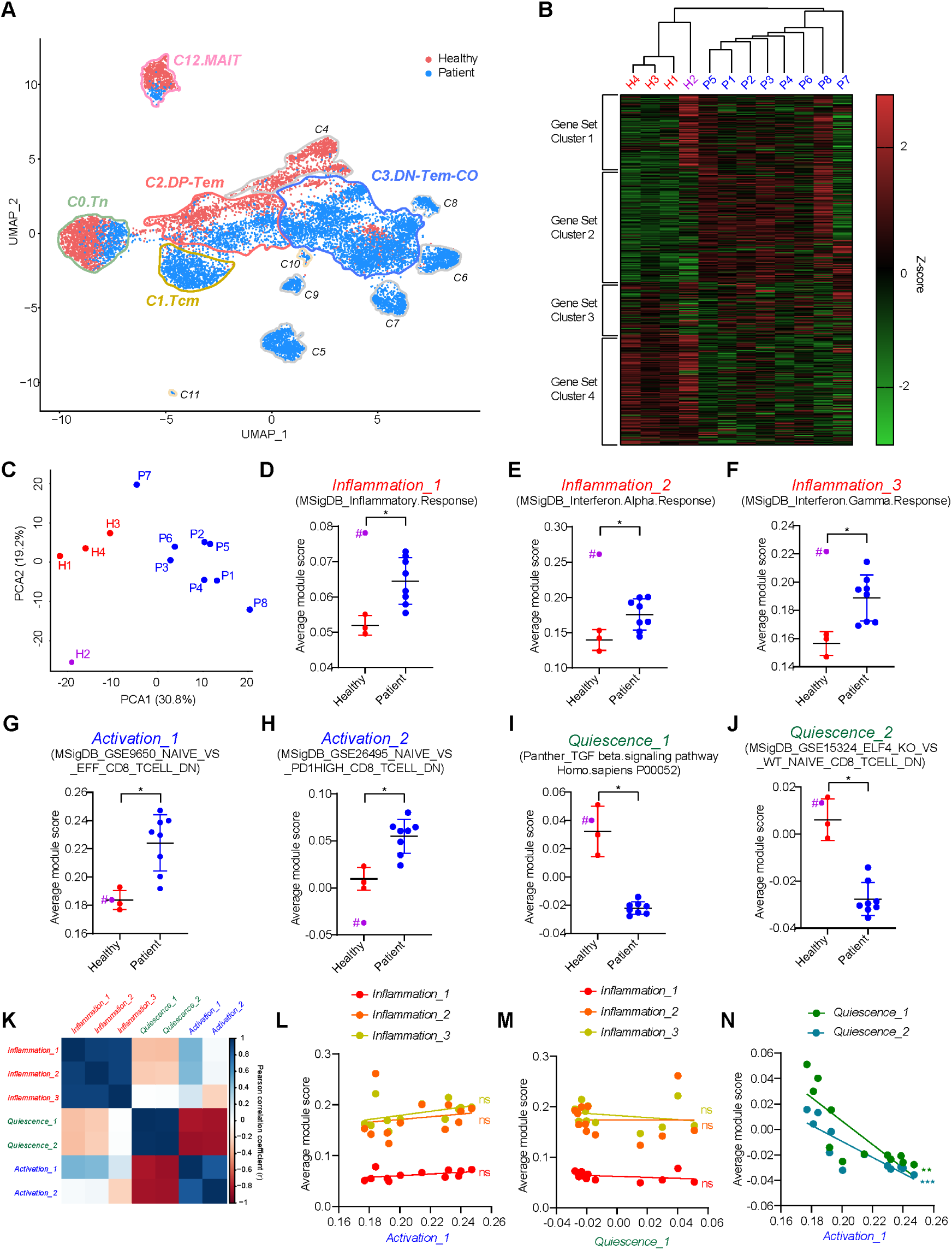
Peripheral blood CD8^+^ T cells in NSCLC patients exhibit altered transcriptional signatures. (**A**) UMAP color coded according to disease status. Colored lines represent boundaries for each cluster. (**B**) Heat map for normalized average module scores of 644 gene sets in *C2.DP-Tem*. Z-score was color-coded from green (low) to red (high). (**C**) PCA plot generated with average module scores of 644 gene sets in *C2.DP-Tem*. (**D**–**J**) Average module scores of indicated gene sets in *C2.DP-Tem*. (**K**) Correlation matrix between gene sets. Pearson correlation coefficient was color coded from red (low) to blue (high). (**L**–**N**) Correlations between the gene sets. Lines represent linear regression. All bar graphs represent mean ± SD. ∗∗∗p<0.001, ∗∗p<0.01, ∗p<0.05. Samples with # were excluded from statistics.

Next, we investigated how the transcriptomic changes are associated with functional alterations. For this, we selected 644 signal and immune-related gene sets from public databases (Broad Institute, PANTHER) which were categorized into four clusters, namely Gene Set Clusters (GSC) 1, 2, 3, and 4, according to the module scores of *C2.DP-Tem* (Fig. 5B). GSC1 and GSC2 showed significant overall increases in patients relative to the levels in healthy individuals, while GSC3 was at a similar level in both groups (Fig. 5B). In contrast, GSC4 was significantly increased in healthy individuals relative to the levels in patients (Fig. 5B). Notably, in the healthy group, one individual (H2), who was excluded from the previous analysis (Fig. 3 and 4), was observed to have a high GSC1 expression level, even higher than all the patients (Fig. 5B), which led to placing H2 in a distinctly different position from the other healthy individuals in the principal component analysis (Fig. 5C).

Next, we analyzed the key functional gene sets within each GSC (Fig. 5, D to J). GSC1 and GSC2, which were significantly higher in patients than in healthy individuals, included gene sets associated with inflammation (*Inflammation,* Fig. 5, D to F) and T cell activation (*Activation*, Fig. 5, G and H), respectively. In contrast, GSC4 contained gene sets related to T cell quiescence(*39-43*) (*Quiescence*), and these were expressed at lower levels in patients than in healthy individuals (Fig. 5, I and J). Interestingly, H2 in healthy individuals showed high *Inflammation* (GSC1); however, *Activation* (GSC2) and *Quiescence* (GSC4) remained unchanged (Fig. 5, D to J, purple). Importantly, in the examination of the expression correlations between these gene sets (Fig. 5, K to N), *Inflammation* did not correlate at all with *Activation* or *Quiescence* (Fig. 5, K to M), whereas *Quiescence* and *Activation* were strongly inversely correlated (Fig. 5, K and N). Changes in these gene sets were not limited to specific cells but were seen across all cells (fig. S4B), suggesting that cancer may systematically and systemically disrupt the function and homeostasis of peripheral blood CD8^+^ T cells.

To determine whether the above phenomenon is cancer-specific or common to diseases involving infection or inflammation, we analyzed published scRNA-seq data from patients with various viral infections and autoimmune diseases (COVID Atlas, Gene Expression Omnibus). COVID-19 patients indeed exhibited higher *Inflammation*, as expected, than healthy individuals (fig. S4, C to E). However, *Activation* and *Quiescence* remained similar to that of healthy individuals (fig. S4, F to I). Remarkably, similar to COVID-19 patients, *Quiescence* was not significantly different from that of healthy individuals among patients with other acute and even chronic viral infections (fig. S4J, yellow box). However, interestingly, various autoimmune diseases showed varying degrees of differences in *Quiescence*, with a particular significant decrease in progressive Sjögren’s syndrome (pSS) (fig. S4J, green box). Furthermore, the decrease in *Quiescence* was not associated with *Inflammation* but was strongly linked to the increase in the *Activation* (fig. S4K). This phenomenon, which was common with several autoimmune diseases and lung cancer, strongly supports the notion that unrestrained T cell quiescence is closely associated with homeostatic dysregulation, resulting in increased T cell activation and/or unregulated differentiation.

### Genetic signatures of homeostatic dysregulation

We further characterized the decreased *Quiescence* and increased *Activation* in terms of gene expression changes. We analyzed the differentially expressed genes (DEGs) between patients and healthy individuals in three key clusters, *C0.Tn, C2.DP-Tem,* and *C3.DN-Tem-CO* (Fig. 6A). Each cluster had 92, 150, and 84 DEGs, respectively, and interestingly, there was a high degree of overlap between these DEGs, with ∼50% of DEGs in each cluster also found in the other clusters (Fig. 6A). Furthermore, the expression patterns of DEGs within each cluster appeared to be regulated in a coordinated manner, as evidenced by the strong positive and negative correlations between their expressions (Fig. 6B and fig. S5, A to C). Based on these observations, we wanted to identify the most significantly changed genes within these coordinated DEGs. Among the shared 19 DEGs between *C0.Tn, C2.DP-Tem,* and *C3.DN-Tem-CO* (Fig. 6, A and C, and fig. S5D), *JUN* and *CISH* were exclusively observed in healthy individuals and patients, respectively, with these two genes being co-expressed in a mutually exclusive manner (Fig. 6C and fig. S5E). Therefore, compared with healthy individuals, patients exhibited significantly elevated *CISH/JUN* ratios (Fig. 6D and fig. S5F). Importantly, the *CISH/JUN* ratio showed a strong correlation with the DEGs (Fig. 6E), indicating that it can function as a genetic marker representing the coordinated gene expression changes observed in patients.

**Fig. 6.**
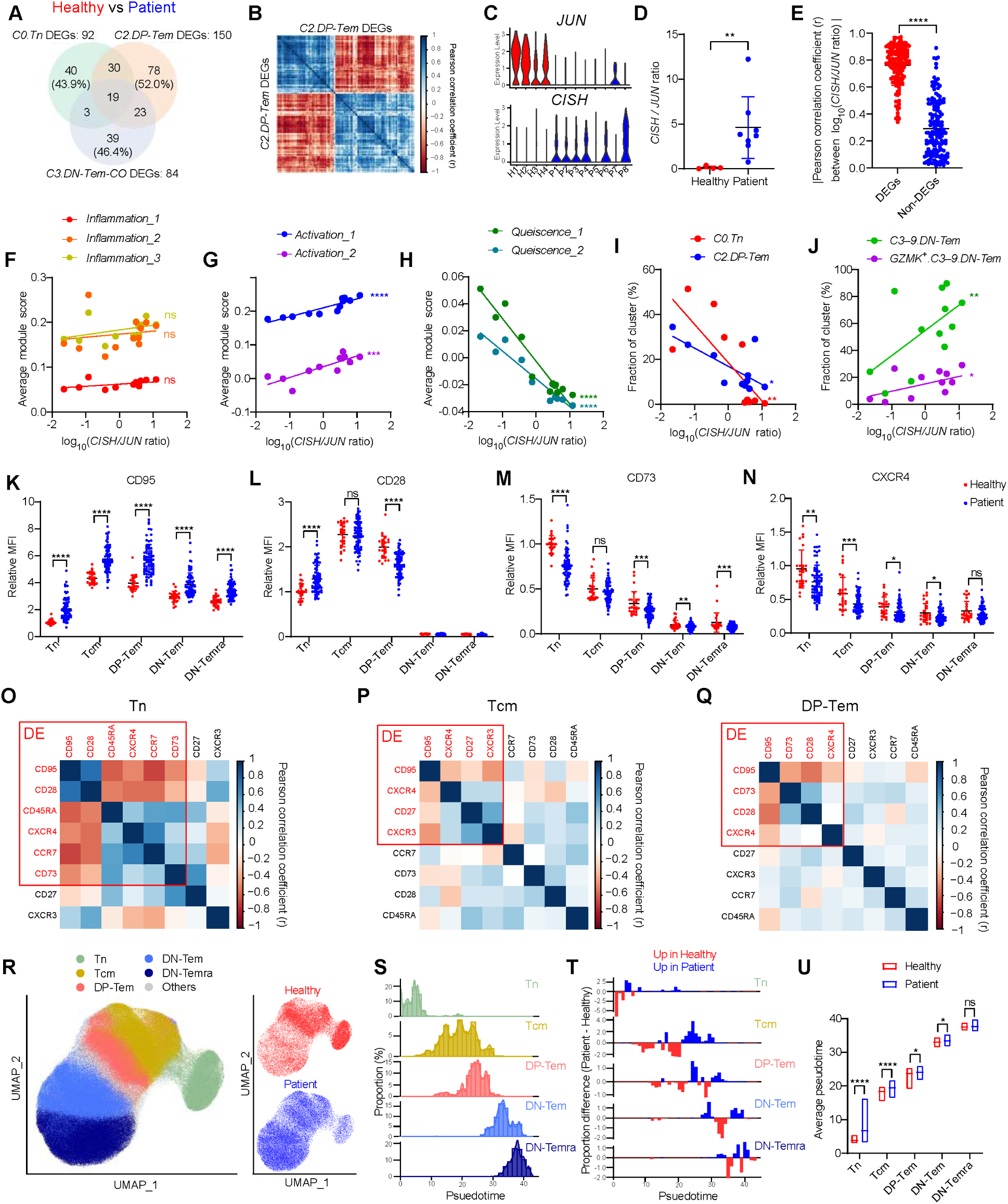
Peripheral blood CD8^+^ T cells in NSCLC patients exhibit coordinated gene and protein expression alterations. (**A**) Venn diagram for DEGs. Numbers in the brackets indicate frequency of unshared DEGs. (**B**) Correlation matrix between 150 DEGs of *C2.DP-Tem*. Pearson correlation coefficient was color coded from red (low) to blue (high). (**C-D**) (C) *JUN* and *CISH* expressions and (D) *CISH/JUN* ratio in *C2.DP-Tem* from healthy individuals (red) and patients (blue). (**E**) Absolute value of Pearson correlation coefficient between *CISH/JUN* ratio and 150 DEGs or randomly selected 150 non-DEGs in *C2.DP-Tem*. (**F**–**J**) Correlation between *CISH/JUN* ratio and (F-H) signature gene sets or (I-J) cluster frequencies. Lines represent linear regression. (**K**–**N**) Relative MFI of indicated molecules in each subset from healthy individuals (n=25) and NSCLC patients (n=71). Relative MFI was calculated by dividing MFIs with that of Tn from a healthy individual. (**O**–**Q**) Correlation matrix between average MFIs of each molecule. Red boxes represent differentially expressed proteins. (**R**) UMAP generated with protein expressions. Cells were color coded according to their labels. (**S**) Histogram of pseudotime in each subset. (**T**) Proportional difference between patient and healthy individuals in each pseudotome point. (**U**) Average pseudotime of each subset in healthy individuals and NSCLC patients. All bar graphs represent mean ± SD. ∗∗∗∗p<0.0001, ∗∗∗p<0.001, ∗∗p<0.01, ∗p<0.05.

Next, we investigated the association between the coordinated gene expression changes (represented by the *CISH/JUN* ratio) and previously observed gene sets (*Inflammation*, *Activation*, *Quiescence*; Fig. 5). Importantly, the *CISH/JUN* ratio was not associated with *Inflammation* at all (Fig. 6F) but was positively correlated with *Activation* (Fig. 6G) and inversely correlated with *Quiescence* (Fig. 6H). Similarly, these changes in the *CISH/JUN* ratio were observed in patients with autoimmune diseases, particularly those with pSS (fig. S5G), in which *Quiescence* decreased the most (fig. S4J). Furthermore, the *CISH/JUN* ratio was strongly correlated with *Quiescence* and *Activation* rather than the *Inflammation* (fig. S5, H to J). More importantly, these phenomena observed at the gene expression levels were also closely associated with differentiation abnormalities previously observed in peripheral blood CD8^+^ T cells—decreases in *C0.Tn* and *C2.DP-Tem* and increases in *C3–9.DN-Tem* and *GZMK^+^.C3–9.DN-Tem* (Fig. 6, I and J). These findings strongly suggest that NSCLC patients undergo coordinated, systemic alterations in the expression of various genes that are associated with dysregulation of peripheral blood CD8^+^ T cell homeostasis.

### Phenotypic signatures of unrestrained T cell activation and differentiation

Based on the above gene expression alterations, we investigated phenotypic markers that indicate the activation and/or differentiation of CD8^+^ T cells in stage IV NSCLC patients. Notably, there were significant differences in the expression levels of various markers (CD95, CD28, CD73, CXCR4, CCR7, CD45RA, CD27, and CXCR3; Fig. 6, K to N and fig. S5, K to N). In particular, CD95 expression, known to increase upon TCR stimulation, was significantly higher in NSCLC patients than in healthy individuals, and surprisingly, this difference was observed across the entire CD8^+^ T cell compartment comprising Tcm, DP- and DN-Tem, DN-Temra and even Tn (Fig. 6K). Furthermore, there was a strong correlation between proteins that are differentially expressed (DE; Fig. 6, O to Q). Similar phenomena were also observed in SCLC patients (fig. S5, O to V). These coordinated protein expression changes appeared to be associated with sequential cell differentiation occurring over time. In fact, when the CD8^+^ T cell compartment was further subclustered by pseudotime analysis based on the above protein expression changes (Fig. 6, R and S), nearly all subsets (especially in Tn, Tcm, and DP Tem) in patients showed enhanced occurrence at the later pseudotimes relative to healthy individuals (Fig. 6, T and U). These results further support the notion that there is a coordinated, systemic alteration in both gene and even protein expression across all peripheral blood CD8^+^ T cell populations in lung cancer patients, leading to a continuous flow of cell differentiation.

### Influence of homeostatic dysregulation on ICI clinical responses

Given the observed gene and protein expression alterations indicative of tumor-associated homeostatic dysregulation, we investigated the potential relationships between these phenomena and the clinical response to ICI therapy. In this regard, we previously reported a strong inverse correlation between CD8^+^ DP-Temra frequency in peripheral blood of NSCLC patients and the abundance of tumor-infiltrating CD8^+^ T cells, with predictive potential for ICI therapy responses(*26*). We therefore hypothesized that a state of low pre-treatment DP-Temra frequency (_pre_DP-Temra^lo^: indicative of high tumor immunogenicity) and low DN-Tem frequency (_pre_DN-Tem^lo^: indicative of high T cell responsiveness) in NSCLC patients would induce the best ICI treatment responses (Fig. 7A). To address this possibility, we retrospectively analyzed _pre_DN-Tem and _pre_DP-Temra frequencies in the peripheral blood of 279 lung cancer patients across four independent cohorts (Fig. 7B).

**Fig. 7.**
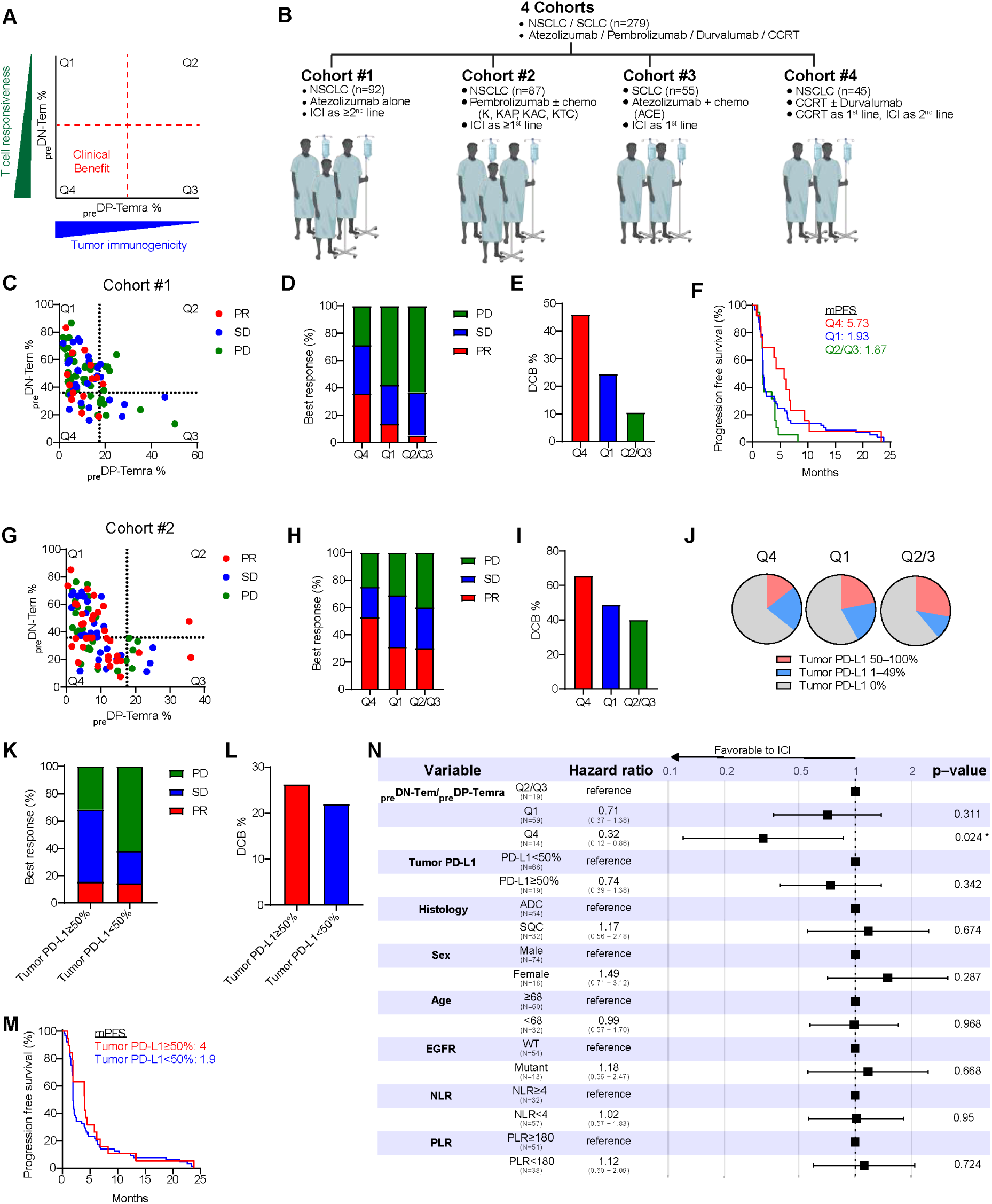
Cancer-associated homeostatic dysregulation in systemic CD8^+^ T cells correlates with ICI responsiveness. (**A**) Strategy to predict clinical benefit using two CD8^+^ T cell subsets. (**B**) Summary of cohorts. (**C**–**F**) Relationship between frequencies of _pre_DN-Tem and _pre_DP-Temra and (C-D) PR, (E) DCB, and (F) PFS in Cohort #1 (n=92). (**G**–**I**) Relationship between frequencies of _pre_DN-Tem and _pre_DP-Temra and (G-H) PR and (I) DCB in Cohort #2 (n=87). (**J**) Tumor PD-L1 expression in Q1–Q4 groups in Cohort #1. (**K**–**M**) Relationship between tumor PD-L1 expression and (K) PR, (L) DCB, and (M) PFS in Cohort #1. (**N**) Forrest plot for hazard ratio between PR and indicated variables in Cohort #1.

In our initial analysis of a cohort of 92 NSCLC patients who received anti-PD-L1 (atezolizumab) monotherapy (Cohort #1; table S4), partial responses (PR) were rarely observed in patients with _pre_DP-Temra^hi^ (>17.2%, Q2 and Q3), whereas it was clearly evident in patients with _pre_DP-Temra^lo^ (<17.2%, Q1 and Q4; Fig. 7, C to D). Particularly, among patients with _pre_DP-Temra^lo^ (Q1 and Q4), those with _pre_DN-Tem^lo^ (<36%, Q4) exhibited higher PR (35.7% vs. 13.6%), durable clinical benefit (DCB; 46.2% vs. 24.6%), and median progression free survival (5.73 months vs. 1.93 months) post-ICI therapy than those with _pre_DN-Tem^hi^ (>36%, Q1) (Fig. 7, D to F). Moreover, this association with clinical responses was also observed in patients receiving anti-PD-1 (pembrolizumab) therapy in combination with chemotherapy (Cohort #2; table S5; Fig. 7, G to I), similar to those receiving anti-PD-L1 (atezolizumab) monotherapy (Fig. 7, C to F). Furthermore, consistent with the findings observed in patients treated with anti-PD-L1 therapy (Fig. 2, K and L), we observed a notable increase in prfn^+^/gnzB^+^ frequency by anti-PD-1 therapy, particularly in patients with _pre_DN-Tem^lo^ (fig. S6A). Importantly, this increase was closely associated with a decrease and an increase in the DP-Tem and DN-Tem frequency, respectively, post-ICI therapy (fig. S6B). Similar trends were observed in SCLC patients who received combined anti-PD-L1 (atezolizumab) and chemotherapy (Cohort #3; table S6; fig. S6, C to E). In contrast, patients who received concurrent chemo-radiation therapy (CCRT; cohort #4), half of whom received anti-PD-1 (durvalumab) consolidation afterwards (table S7 and fig. S6F), showed no difference in clinical outcomes between Q4 and Q1 (fig. S6, G and H); however, the clinical benefit of anti-PD-1 consolidation was observed exclusively in Q4 but not Q1 (fig. S6, I to K). These data from lung cancer patients strongly support an immunological link between these two events, namely, altered DN-Tem (and DP-Temra) frequency and therapeutic responses to ICI.

We next investigated whether the predicted difference in ICI responses based on _pre_DN-Tem/_pre_DP-Temra was associated with tumor PD-L1 expression widely used in clinics. However, the higher ICI response rates of Q4 patients were not associated with tumor PD-L1 expression (Fig. 7J), and indeed, within Cohort #1, tumor PD-L1 expression did not affect ICI therapy responses (PR, DCB, and mPFS; Fig. 7, K to M). Furthermore, when compared with various clinicopathologic variables, no clinical information other than _pre_DN-Tem was significantly associated with ICI therapy responses (Fig. 7N). Given the patient-specific *GZMK^+^.DN-Tem* accumulation (Fig. 4), we further explored the relationship between *GZMK^+^.DN-Tem* frequency and ICI therapy responses. Based on the higher expression of *CXCR3* in *GZMK^+^.DN-Tem* than in *GZMK^−^.DN-Tem* (fig. S6, L and M), the increased proportion of CXCR3^+^.DN-Tem in patients (fig. S6N), and the relatively early pseudotime of CXCR3^+^.DN-Tem (fig. S6O), we used CXCR3^+^.DN-Tem (akin to *GZMK^+^.DN-Tem*) for predicting ICI therapy responses. As expected, among the patients with _pre_DP-Temra^lo^, those with low CXCR3^+^.DN-Tem frequency exhibited the best outcomes (fig. S6, P to R). Collectively, these results suggest that the cancer-associated systemic alterations in peripheral blood CD8^+^ T cell populations can significantly influence clinical responses to ICI therapy.

## DISCUSSION

The immunity cycle between tumors and antigen-specific CD8^+^ T cells is a major barrier against the malignant transformation of cancer(*44, 45*). However, tumors also adapt various resistance mechanisms against CD8^+^ T cells, including engagement of immune checkpoints(*33, 34, 46, 47*). Although a state of exhaustion is known to inhibit tumor-reactive CD8^+^ T cells within tumor tissues(*48-50*), how tumors disrupt systemic CD8^+^ T cell populations to promote malignancy is less understood. In this study with lung cancer patients, we demonstrated homeostatic dysregulation of peripheral blood CD8^+^ T cells with loss of quiescence and increased spontaneous activation/differentiation. This process occurred systemically and progressively during tumor progression, led to gradual accumulation of DN-Tem that are refractory to TCR-driven proliferation, and moreover, strongly correlated with poor clinical responses to ICI therapy. Therefore, these findings suggest a new cancer evasion mechanism that could potentially avoid tumor-reactive CD8^+^ T cells at the systemic level.

The most notable feature seen in our patient cohort was the remarkable increase in the DN-Tem frequency. It is important to note that differentiation into DN-Tem is a normal immune-related process, as this subset is commonly observed in the blood of healthy individuals with a proportional increase with age(*51-53*). Thus, while it was not surprising to see DN-Tem in the patients, the accelerated rate of accumulation was clearly abnormal and of great interest. Given the high expression of cytotoxic molecules (prfn/gnzB) in DN-Tem, we assumed that tumor-associated, chronic antigenic stimulation would primarily contribute to this acceleration. However, since most, if not all, DN-Tem are likely to be tumor non-specific bystander cells(*30, 31, 54-56*), we also postulated a process involving antigen-independent DN-Tem accumulation. Indeed, our scRNA-seq data provided strong evidence that these two possibilities occur simultaneously and are equally important. As expected, increased clonality in DN-Tem was evident in patients, clearly supporting a role for oligoclonal expansion. However, the most surprising and unexpected data was that despite the increase in clonality, TCR diversity of DN-Tem did not decrease due to an unexpected increase in the number of unique clonotypes with little expansion. Therefore, the accelerated accumulation of DN-Tem (and conversely, the decrease of DP-Tem) in patients is thought to occur, at least in part, through a continuous proliferation-independent differentiation from DP-Tem to DN-Tem during tumor progression.

The unexpected findings mentioned above raised the question of the possible complexity in the properties and composition of DN-Tem. Further dissection of this subset revealed that two major subsets could be separated by *GZMK* expression, namely *GZMK^−^.DN-Tem* and *GZMK^+^.DN-Tem*. Among them, *GZMK^−^.DN-Tem-BEC/MEC* (<10 clones) was thought to reflect its long history of an antigen-specific response, and its appearance in both patients and healthy individuals suggested that this subset was commonly generated via typical immune responses before tumor onset (fig. S7A). However, relatively low diversity and high clonality of *GZMK^−^.DN-Tem-HEC* (≥10 clones), which was observed only in patients, suggested that this subset had undergone massive proliferation, presumably to tumor antigens (fig. S7A). Therefore, it seems clear that *GZMK^−^.DN-Tem* played a role in accelerating the accumulation of DN-Tem in patients via robust clonal expansion.

In contrast to its role in increasing clonality, *GZMK^−^.DN-Tem* showed a negligible contribution to TCR diversity, which we described above as an unexpected finding. We demonstrated here that unlike *GZMK^−^.DN-Tem*, *GZMK^+^.DN-Tem* contributed significantly to shaping the relatively high TCR diversity of DN-Tem. Although *GZMK^+^.DN-Tem* can be considered as a transient subset, we noticed the increase in *GZMK^+^.DN-Tem-BEC/MEC* was not accompanied by a subsequent increase in *GZMK^−^.DN-Tem-BEC/MEC* and notably that some *GZMK^+^.DN-Tem* contained tumor antigen-irrelevant, viral-specific clones. Therefore, we believe that *GZMK^+^.DN-Tem* in patients is not a transient subset appearing during differentiation into *GZMK^−^.DN-Tem* but rather a unique discrete subset generated by diverse clonotypes (fig. S7A).

The above findings, that *GZMK^+^.DN-Tem* abundance was greater in patients than in healthy individuals and had tumor non-specific bystander (viral-specific) clones raised an intriguing question as to its generation mechanisms. Initially, we thought that cancer-associated chronic inflammation, which is common in malignant diseases(*57-60*), could be a key factor in accelerating the DP-to-DN transition and promoting the generation of *GZMK^+^.DN-Tem*. In fact, our pathway analysis revealed significantly increased inflammation signatures in lung cancer patients. These signatures were generally lower in healthy people, but were unusually high in one healthy individual (H2). Nevertheless, the situation of H2 with high inflammation signatures was very different from that of patients; while all patients showed high levels of T cell activation signatures, H2 showed low levels, similar to other healthy individuals. A similar phenomenon seen in H2 was also observed in COVID-19 patients. Likewise, patients with various viral infections and even autoimmune diseases showed no correlation between inflammation and T cell activation signatures, which was consistent with our data from lung cancer patients. We therefore thought that different mechanisms other than cancer-associated chronic inflammation might be involved.

Unlike the inflammation signatures, we observed a strong association between the T cell quiescence signatures and all aberrant changes in systemic CD8^+^ T cell compartment, including accelerated DN-Tem accumulation. Interestingly, peripheral blood CD8^+^ T cells from lung cancer patients showed markedly reduced quiescence signatures with a strong inverse correlation with activation signatures. We also confirmed that such low quiescence and high activation signatures were closely linked to an increase in DN-Tem (and *GZMK^+^.DN-Tem)* suggesting that there were significant alterations in the ability to maintain systemic CD8^+^ T cells in a quiescent state during tumor progression. This notion was further supported by our data showing coordinated alterations in the expression of various genes and proteins, indicative of T cell activation and differentiation. All these data highlight the role of cancer-associated homeostatic dysregulation in promoting uncontrolled activation/differentiation of systemic CD8^+^ T cell populations (fig. S7B).

The above findings leave open question of what stimuli can trigger this phenomenon and, importantly, whether it is limited to lung cancer patients only. In this study, we demonstrated that the loss of T cell quiescence seen in lung cancer patients was not commonly observed in acute and chronic viral infections. However, to our surprise, we noticed that in some autoimmune diseases, especially in patients with pSS, T cell quiescence signatures were significantly decreased with inverse correlation with T cell activation signatures. It is unclear whether this association is just coincidental, but it is worth mentioning that pSS patients were reported to have a relatively high risk of developing various cancers, including lung cancer and lymphoma(*61-63*).

Since DN-Tem from healthy individuals and patients showed high levels of prfn/gnzB, we assume that if there are tumor-reactive cells within this subset, they are fully functional to exert potent cytotoxic activity. However, we showed that DN-Tem was almost incapable of proliferating upon TCR stimulation *in vitro*, and was also poorly responsive to proliferate after ICI therapy. Therefore, rather than disrupting the cytotoxic function of DN-Tem *per se*, cancer is thought to induce a persistent homeostatic perturbation that switch cells from high division potentials (DP-Tem) to low division potentials (DN-Tem) in response to antigenic stimulation. This notion is reinforced by our intensive retrospective analysis data from four independent cohorts of lung cancer patients receiving anti-PD-(L)1 therapy, which showed that _pre_DN-Tem^hi^ patients exhibited poorer clinical responses compared to _pre_DN-Tem^lo^ patients. As a strong proliferative response of CD8^+^ T cells in the blood has been shown to be important for response outcomes after ICI(*64-67*), we believe that the poor ICI responses in _pre_DN-Tem^hi^ patients were primarily due to the inherently weak proliferative potential of DN-Tem.

Building on the above relationship between _pre_DN-Tem frequency and post-ICI response, we propose a new mechanism by which cancer evades attack of potentially tumor-reactive CD8^+^ T cells. As such, cancer induces homeostatic dysregulation with loss of T cell quiescence and uncontrolled activation/differentiation across the entire CD8^+^ T cell populations. This is followed by a progressive accumulation of DN-Tem. Despite its high cytotoxic activity, DN-Tem has a low proliferative potential upon antigenic TCR stimulation, therefore causing a gradual shift in the fitness of potentially tumor-specific CD8^+^ T cells to a non-reactive state. We now named this phenomenon “Cancer-associated Homeostatic dysregulation Accelerating uncOntrolled differentiation of Systemic CD8^+^ T cells (CHAOS)” (fig. S7C). Under the condition of CHAOS, we suggest that even cancer patients who could have responded to ICI therapy will eventually accumulate DN-Tem and become non-responders. Therefore, it will be important to explore the possibility that patients with stage IV cancer who are _pre_DN-Tem^hi^—presumably ICI non-responders based on CHAOS—respond much better to alternative therapies other than ICI, as demonstrated in our control cohort of lung cancer patients receiving CCRT. In conclusion, we uncover a new mechanism of cancer immune evasion that has important clinical implications for improving our understanding of the interactions between cancer and CD8^+^ T cells and for developing effective therapeutic strategies for cancer patients.

## Acknowledgments

We thank professors Deok Hwan Yang, Sang Yun Song, Ik Joo Chung, and Joon Haeng Rhee from CNU for helpful advice and clinical comment. We also thank the Biobank of CNU Hwasun Hospital for providing biospecimens used for this study. Some of the figures were created using Biorender.com.

## Funding

This study was supported by:

Korean Ministry of Science and ICT grant 2020R1A5A2031185 (JHC)

Korean Ministry of Science and ICT grant 2020M3A9G3080281 (JHC)

Korean Ministry of Science and ICT grant 2022R1A2C2009385 (JHC)

Korean Ministry of Education grant 2022R1A6A3A01086438 (SWL)

Korean Ministry of SMEs and Startups Collabo R&D project RS-2023-00221640 (JHC)

## Author contributions

Conceptualization: SWL/JHC Methodology: SWL/JHC Software: SWL/JHC

Formal Analysis: SWL/JHC Investigation: SWL/IJO/JHC

Resources: SWL/JSY/YJK/HJC/CKP/IJO/JHC Data Curation: SWL/CKP/IJO/JHC

Writing-Original Draft: SWL/JHC

Writing-Review&Editing: SWL/HOK/CKP/IJO/JHC Visualization: SWL/JHC

Supervision: IJO/JHC

Project Administration: IJO/JHC Funding Acquisition: SWL/JHC

## Data and materials availability

All data are available in the main text or the supplementary materials. Single-cell RNA-seq data has been deposited at GEO (GSE247754) and are publicly available as of the date of publication.

## Supplementary Materials

Materials and Methods

Figs. S1 to S7

Tables S1 to S7

