## Supplemental Materials for "Homeostatic Dysregulation of Systemic CD8^+^ T Cell Compartment in Lung Cancer Patients"

**The PDF file includes:**

Materials and Methods
Figs. S1 to S7
Tables S1 to S7

### Materials and Methods

#### Human samples

Peripheral blood of the patients was provided by the Biobank of Chonnam National University Hwasun Hospital, a member of the Korea Biobank Network. Information on the analyzed patients is summarized in table S1. We enrolled patients with histologically or cytologically confirmed lung cancer and divided them into 4 cohorts as follows: stage IV non-small cell lung cancer (NSCLC) who received immune checkpoint inhibitors (ICIs) as the second- and later-line (Cohort #1) or the first-line treatment (Cohort #2), extensive disease (ED) stage small cell lung cancer (SCLC) who received ICI plus chemotherapy as the first-line treatment (Cohort #3), and stage III NSCLC who received concurrent chemoradiation therapy (CCRT) followed by ICI consolidation treatment (Cohort #4). All patients provided written informed consent in accordance with local regulations (South Korea) and with institutional review board (IRB) approval. Pretreatment peripheral blood was collected 1−10 days before or on the initial date of ICI treatment (Cohorts #1, #2, and #3) or CCRT (Cohort #4). ICIs were administered intravenously for 60 min every 3 weeks (Cohorts #1, #2, and #3). For patients who completed CCRT without progression (Cohort #4), ICI (durvalumab) was administered every 2 weeks. For patients with Cohort #1 to #3, post-treatment blood samples were collected 7−14 days after the first ICI treatment. ICI treatment continued until confirmed progression, initiation of alternative cancer therapy, unacceptable toxicity, or the occurrence of other reasons to discontinue the drug. In Cohort #4, durvalumab consolidation treatment continued for a maximum duration of 12 months until the aforementioned events. Most of the blood from healthy individuals was acquired from the Korean Red Cross (Blood donors). Some of the healthy individuals were recruited at Chonnam National University Hwasun Hospital. The information about blood donors from the Korean Red Cross was not accessible due to local regulations. Peripheral whole blood was collected from the cephalic vein using BD Vacutainer (BD) and immediately processed with Lymphoprep (Stemcell technologies) to obtain peripheral blood mononuclear cells (PBMCs). PBMCs were fast frozen at −80ºC using 10% dimethyl sulfoxide (Merck) in fetal bovine serum (FBS) (Gibco), then thawed in a 37ºC water bath upon experiment.

#### Flow cytometry and sorting

Thawed PBMCs were washed twice with complete media (RPMI-1640 (Welgene) supplemented with 10% FBS, Penicillin-Streptomycin (Welgene), 2-mercaptoethanol (Welgene), non-essential amino acid (Welgene), glutamine (Welgene), and HEPES (Welgene)), and used for further experiments. Surface molecules were stained in staining media (PBS (Welgene) supplemented with 3% FBS and EDTA (Bioneer)) for 30 min. For intracellular molecules, samples were fixed and permeabilized with BD Cytofix/Cytoperm (BD) or FOXP3/Transcription Factor Staining Buffer Set (eBioscience) for 20 min, then stained for intracellular molecules for 30 min. All processes were done on ice. Prepared samples were run using CytoFLEX S (Beckman Coulter), or CytoFLEX LX (Beckman Coulter). For fluorescence-activated cell sorting (FACS), prepared samples were run using CytoFLEX SRT (Beckman Coulter). Data were analyzed using Flowjo software (Tree Star) and visualized using Prism (GraphPad). The frequencies of Tn, Tcm, Tem, and Temra were calculated relative to $CD8^{+}CD3^{+}$ cells. The frequencies of DP ($CD27^{+}CD28^{+}$), SP ($CD27^{+}CD28^{-}$ and $CD27^{-}CD28^{+}$), DN ($CD27^{-}CD28^{-}$), and $CXCR3^{+}$.DN ($CXCR3^{+}CD27^{-}CD28^{-}$) phenotypes were calculated relative to $CD8^{+}CD3^{+}CCR7^{-}CD45RA^{-}$ for Tem subsets or $CD8^{+}CD3^{+}CCR7^{-}CD45RA^{+}$ for Temra subsets. Following antibodies for flow cytometry were purchased from Biolegend, BD, or Invitrogen: anti-CD8α (RPA-T8), anti-CD4

(A161A1), anti-CD3 (OKT3), anti-CCR7 (G043H7), anti-CD45RA (HI100), anti-CD27 (LG.7F9), anti-CD28 (CD28.2), anti-IL2 (MQ1-17H12), anti-CD127 (hIL-7R-M21), anti-CD45RO (UCHL1), anti-CXCR3 (G025H7), anti-TCF1 (7F11A10), anti-CD44 (BJ18), anti-CD69 (FN50), anti-PD-1 (J105), anti-CD95 (DX2), anti-CD103 (Ber-ACT8), anti-ICOS (C398.4A), anti-CD5 (L17F12), anti-KLRG1 (14C2A07), anti-CD56 (5.1H11), anti-Bcl2 (100), anti-CD122 (TU27), anti-Ki-67 (Ki67), anti-CD62L (DREG-56), anti-CD40L (24-31), anti-IFN-γ (B27), anti-CD39 (A1), anti-Perforin (B-D48), anti-Granzyme B (GB11), anti-CD57 (HCD57), anti-CXCR4 (12G5), and anti-CD73 (AD2). In some experiments, fluorochrome-conjugated monoclonal antibodies (for CD8, CD27, and CD45RA) generated by SELECXINE were used.

**Patient grouping strategies**

To group patients according to perforin (prfn) fold change (FC) and granzyme B (gnzB) FC, FCs were acquired by dividing prfn$^+$ (and gnzB$^+$) frequencies after ICI therapy with baseline prfn$^+$ (and gnzB$^+$) frequencies of the respective patient. Patients with high FC (FC≥1.1; ~30% of total patients) were considered FC$^{hi}$ and the rest as FC$^{lo}$. To group patients according to baseline prfn, gznB, DN-Tem, or DP-Tem, frequencies of these cells in total Tem were accessed. Top ~30% were considered patients with high frequency of these cells. To investigate biomarker efficiency of DN-Tem and DP-Temra, DN-Tem and DP-Temra frequencies were assessed in total Tem and total Temra, respectively, in peripheral blood of lung cancer patients and healthy individuals. Patients with DN-Tem and DP-Temra frequencies higher than that of most (>~80%) healthy individuals who were analyzed under the same experiment settings (DN-Tem≥36% and DP-Temra≥17.2% for NSCLC patients and DN-Tem≥36% and DP-Temra≥11% for SCLC patients) were considered DN-Tem$^{hi}$ and DP-Temra$^{hi}$. The threshold for CXCR3$^+$.DN-Tem (≥1.25%) was determined with a similar approach.

***In vitro* T cell proliferation**

Anti-CD3 (clone OKT3; 5μg/ml) and anti-CD28 (clone CD28.2; 2μg/ml) antibodies were coated onto a flat-bottom 96-well immuno-plate (Thermo Fisher Scientific) overnight at 4ºC. DP-Tem and DN-Tem were FACS-purified, then labeled with CellTrace$^{TM}$ Violet (CTV; Thermo Fisher Scientific). The labeled cells were seeded onto the coated plate ($10^4$ cells/well) and cultured for 5 days in a 37 ºC $CO_2$ incubator (DAIHAN Scientific). CTV-dilution was assessed with flow cytometry.

**Single cell RNA sequencing**

Cells were resuspended in PBS and filtered through a 40 μm filter. After cells were counted using LUNA-FL™ Automated Fluorescence Cell Counter (Logos Biosystems), 10X Genomics Chromium Instrument and cDNA synthesis kit (Chromium Next GEM Single Cell 5′ Kit v2 & Chromium Next GEM Chip K Single Cell Kit) were used to generate a barcoded cDNA library for single cell RNA-sequencing. cDNA library quality was determined using an Agilent Bioanalyzer (Agilent technologies). Using this library, two paired-end 200bp FlowCells were run on an Illumina NovaSeq6000 S2 Rgt Kit v1.5 (200 cycles (Read lengths: 28 bp Read1, 10 bp I7 Index, 10 bp I5 Index, and 90 bp Read2) (Illumina). For T cell receptor (TCR) sequencing, cDNA was used to process a nested-PCR enrichment method in order to increase specificity of the amplification of the constant region of the T-cell transcripts. Using 10X Genomics Chromium Single Cell Human TCR Amplification Kit, TCR libraries were enriched for Alpha-beta (TRA/B) TCRs. TCR library quality was determined using an Agilent Bioanalyzer (Agilent technologies). Using this library, two paired-end 200bp FlowCells were run on an Illumina NovaSeq6000 S2 Rgt Kit v1.5 (200 cycles (Read lengths: 50 bp Read1, 10 bp I7 Index, 10 bp I5 Index, and 100 bp Read2) (Illumina). All scRNA-seq analysis were carried out using Seurat R package. De-multiplexed filtered gene barcode matrices were loaded and merged. The merged matrix was processed with quality-control, normalization, and scaling. Next, variable features were used for principal component analysis (PCA), which was then used for UMAP. CITE-seq was merged as a new assay and TCR information was merged as metadata for further analysis.

**Bioinformatics**

Clusters were defined using shared nearest neighbor modularity optimization-based clustering algorithm. Their clonality and clonal diversity were calculated using Pielou's evenness index (asbio R package) and Shannon diversity index (diverse R package), respectively. Clonotypes were defined using complementarity-determining regions (CDRs) of TCRα and TCRβ chains acquired from single cell TCR sequencing. The number of unique clonotypes was assessed by counting duplicated clonotypes as one. The number of duplicates of each unique clonotype was counted, and the clonotypes were categorized as BEC (no duplicates), MEC (2–9 duplicates), or HEC ($\geq$ 10 duplicates) based on the number of duplicates. The virus-specific clonotypes were determined based on the CDR3 sequences of the TCRα and TCRβ chains, which were considered virus-specific if they both matched the same viral epitope-specific CDR3 sequences specified in VDJdb. Pseudotime analysis of scRNA-seq data was carried out using monocle3 R package. Seurat data was transformed into a monocle3-compatible format using SeuratWrappers R package. The transformed data was clustered using the Louvain method in the UMAP. *C0.Tn* cluster was selected as the beginning of the pseudotime. The calculated pseudotime was then merged as metadata to the former Seurat data and used for further analysis. For module score analysis, module scores for each gene set were calculated in total $CD8^+$ T cells using AddModuleScore function in Seurat R package. The average module score of the *DP-Tem* cluster in each individual was calculated and used for further analysis. For some experiments, the following published scRNA-seq data sets from peripheral blood of healthy individuals and patients with viral infections or autoimmune diseases were used: COVID ATLAS (COVID19), GSE149689 (influenza virus), GSE202410 (HIV), GSE182159 (HBV), GSE125527 (systemic lupus erythematosus), GSE152316 (Crohn's disease and spondyloarthritis), GSE138266 (multiple sclerosis), GSE125527 (ulcerative colitis), GSE185857 (palmoplantar pustulosis), and GSE157278 (progressive Sjögren's syndrome). Clusters with $CD8^+$ DP-Tem signatures (*GZMH*$^-$*GZMK*$^+$) in these datasets were accessed and used for module score analysis. Relative module scores were calculated by subtracting each patient's average module score with the average of that of healthy individuals from the same dataset. Differentially expressed genes (DEGs) were determined using the Wilcoxon Rank Sum test adjusted with Bonferroni correction. Genes that had significant p-values and high expression fold changes (adjusted p-value$\leq$0.05 and |logFC|$\geq$0.3) were considered DEGs. The expression correlation between DEGs was calculated by accessing the average expression of each gene for each individual and then finding the Pearson correlation coefficient between them. *CISH/JUN* ratio was calculated by dividing the average *CISH* expression with the average *JUN* expression of *DP-Tem* (*GZMH*$^-$*GZMK*$^+$) cluster in each individual. The relative *CISH/JUN* ratio was calculated by subtracting each patient's $\log_{10}$(*CISH/JUN* ratio) with that of healthy individuals from the same dataset.

**Quantitative PCR**
Total RNAs were collected from total PBMCs ($10^6$ cells) using Hybrid-R$^{TM}$ (GeneAll) and then used to synthesize cDNA with TOPscript cDNA synthesis Kit (enzynomics). The cDNAs were then used for quantitative PCR using PowerUp$^{TM}$ SYBR$^{TM}$ Green Master Mix (applied biosystems) with the following primers on a CFX Opus 384 (Bio-Rad):
JUN-forward: TCCAAGTGCCGAAAAAGGAAG
JUN-reverse: CGAGTTCTGAGCTTTCAAGGT
CISH-forward: TTCGGGAATCTGGCTGGTATT
CISH-reverse: GAACGTGCCTTCTGGCATCT

**Protein expression analysis**
To analyze protein expression changes between healthy individuals and patients, we accessed MFIs of each protein using flow cytometry. To normalize for batch effects across experiments, a healthy individual was included in all experiments and served as the reference for analysis. For all proteins, the MFI of each subset was divided by the MFI of Tn of the reference healthy individual to get the relative MFI. Differentially expressed (DE) proteins were determined using p-values by Student's T-test and fold changes (p-value $<0.05$ and FC$>1.1$ or FC$<0.909$). The expression correlation between proteins was calculated by accessing the average relative expression of each protein for each individual and then finding the Pearson correlation coefficient between them. For pseudotime analysis with proteins, the following molecules were analyzed using flow cytometry: CD8, CD3, CXCR4, CD45RA, CD95, CD27, CD28, CD73, CCR7, and CXCR3. Channel values of flow cytometric data were accessed and analyzed with Seurat R package. The Seurat data was scaled but not normalized, and analyzed for UMAP. Pseudotime analysis was performed using monocle3 R package. Each $CD8^+$ T cell subset was divided into one pseudotime interval and the pseudotime distribution for each subset in healthy individuals and patients was accessed. For each pseudotime interval, the distribution of the patients' subset was subtracted by the distribution of healthy individuals' subset at that pseudotime to obtain the proportion difference. The average pseudotime was calculated for each individual in the indicated subsets and then used for comparison.

**Clinical analysis**
During ICI treatment, a follow-up CT scan for the response assessment was performed every 2 to 3 cycles (Cohort #1, #2, #3) or 4 to 6 cycles (Cohort #4) of ICI treatment. The clinical response to the treatment was defined by the Response Evaluation Criteria in Solid Tumors (RECIST) version 1.1. The best response was defined as complete response (CR) if cancer was no longer detectable, partial response (PR) if the size of the cancer decreased, stable disease (SD) if there was no progression, and progressive disease (PD) if the size of the cancer increased on CT up to 3 months after starting the treatment. In addition, PR or SD by 6 months was defined as durable clinical benefit (DCB). Forrest plot was analyzed using the indicated variables against PR. Hazard ratios were calculated against one factor (labeled as reference) in each variable.

**Ethics approval**
This study was approved by the Institutional Review Boards of Chonnam National University Medical School and Hwasun Hospital (CNUHH-2018-036 and CNUHH-2021-045). All patients provided written informed consent.

**Statistics**

Statistical analysis was performed with Prism (GraphPad) or R. The statistical significances were tested with Mann-Whitney U test for unpaired samples or Wilcoxon matched-pairs for paired samples unless addressed otherwise. Linear regression was performed using Prism, and p-values were calculated using an F-test for non-zero correlation. Values of ∗∗∗∗$p<0.0001$, ∗∗∗$p<0.001$, ∗∗$p<0.01$, ∗$p<0.05$ were considered significant.

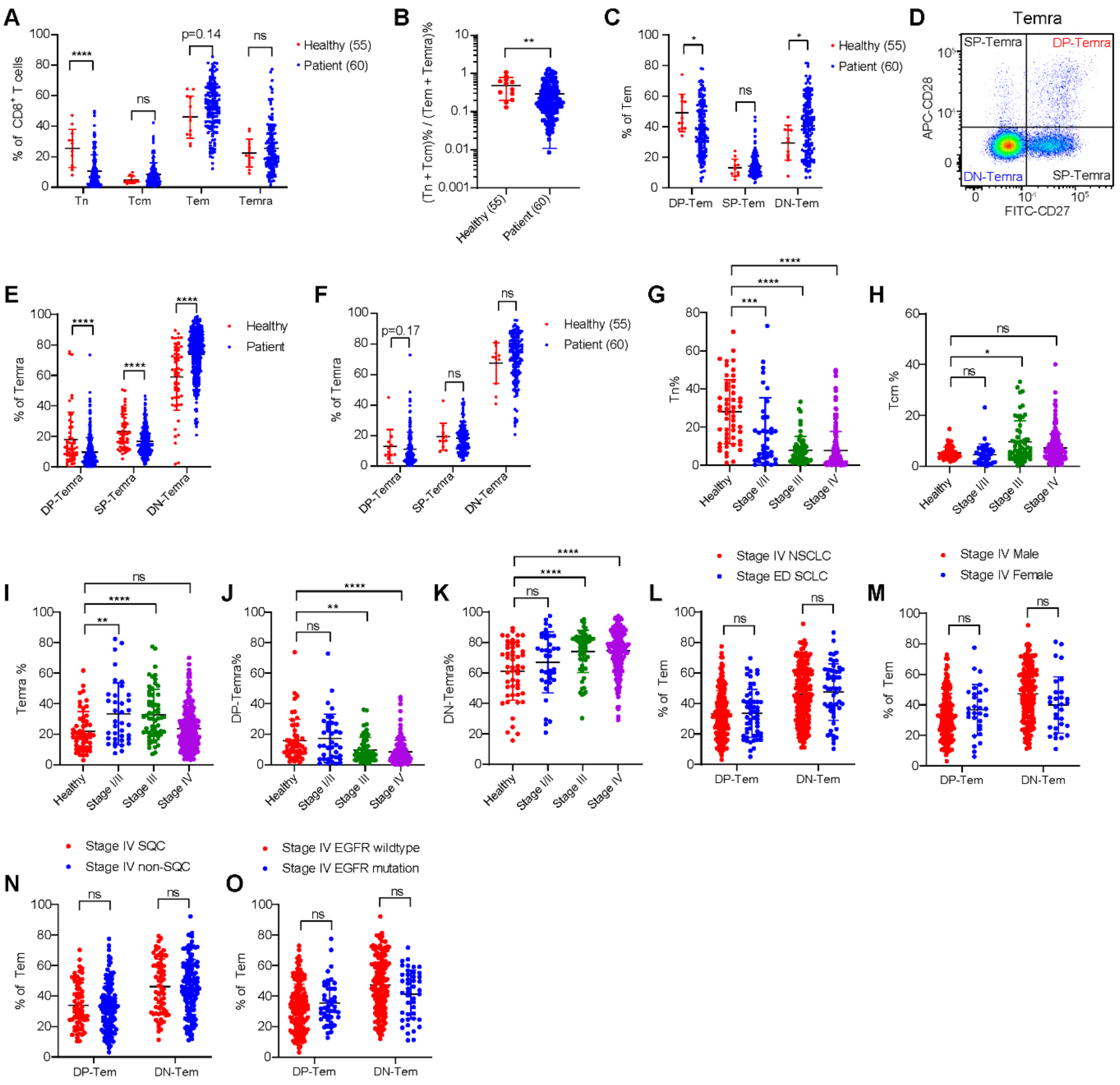


**Fig. S1. Lung cancer patients exhibit stage-dependent accumulation of a more-differentiated subset of peripheral blood CD8$^+$ Tem.**

(**A**–**C**) Age-matched comparisons for (A) CD8$^+$ T cell subset frequencies, (B) ratio between less-differentiated (Tn and Tcm) and more-differentiated (Tem and Temra) subsets, and (C) CD8$^+$ Tem subset frequencies in healthy individuals (n=12) and patients (n=144). Individuals with age between 40 to 70 were selected. Numbers in brackets represent mean age of the group.

(**D**) Gating strategy for CD8$^+$ Temra subsets.

(**E**–**F**) Temra subset frequencies (E) in healthy individuals (n=53) and patients (n=349), (F) and in age-matched healthy individuals (n=12) and patients (n=144).

(**G**–**K**) Frequencies of (G) Tn, (H) Tcm, (I) Temra, (J) DP-Temra, and (K) DN-Temra in patients at different stages of non-small cell lung cancer (NSCLC) (n=53, 37, 60, and 197, respectively).

(**L**) DP-Tem and DN-Tem frequencies in stage IV NSCLC patients or in extensive disease (ED) stage small cell lung cancer (SCLC) patients (n=197 and 55, respectively).

(**M–O**) $CD27^{+}CD28^{+}$ double positive (DP)-Tem and $CD27^{-}CD28^{-}$ double negative (DN)-Tem frequencies in stage IV NSCLC patients grouped by (M) sex (n=167 and 30), (N) histological classification (n=65 and 132), and (O) epidermal growth factor receptor (EGFR) mutation (n=153 and 44).
All bar graphs represent mean ± SD, ∗∗∗∗p<0.0001, ∗∗∗p<0.001, ∗∗p<0.01, ∗p<0.05.

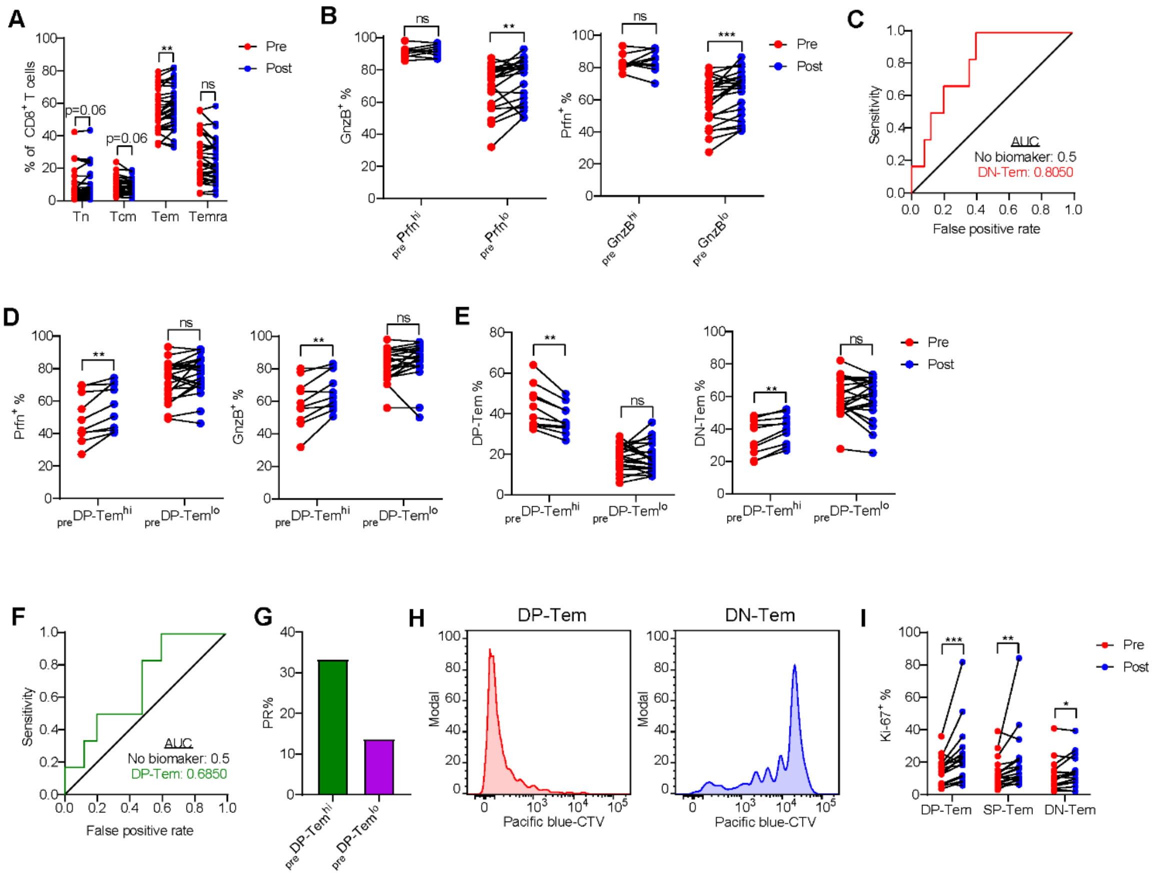


**Fig. S2. Decreased frequency of DP-Tem correlates with poor clinical responses to ICI therapy.**
(**A**) Changes in $CD8^+$ T cell subsets after immune checkpoint inhibitor (ICI) therapy (n=32).
(**B**) Changes in granzyme $(gnzB)^+$ (or perforin $(prfn)^+$) frequencies after ICI therapy in $_{pre}Prfn^{hi}$ (n=9) and $_{pre}Prfn^{lo}$ (n=23) groups (or in $_{pre}GnzB^{hi}$ (n=9) and $_{pre}GnzB^{lo}$ (n=23) groups).
(**C**) Receiver operating characteristic (ROC) curve of DN-Tem (from DN-$Tem^{lo}$ to DN-$Tem^{hi}$) and its area under curve (AUC).
(**D–E**) Changes in (D) $prfn^+$ and $gnzB^+$ frequencies and (E) DP-Tem and DN-Tem frequencies after ICI therapy in $_{pre}$DP-$Tem^{hi}$ and $_{pre}$DP-$Tem^{lo}$ groups (n=10 and 22, respectively).
(**F**) ROC curve of DP-Tem (from DP-$Tem^{hi}$ to DP-$Tem^{lo}$) and its AUC.
(**G**) Proportion of patients with partial response (PR) to ICI therapy.
(**H**) CTV-dilution of DP-Tem and DN-Tem after 5 days of *in vitro* activation with plate-bound anti-CD3 and anti-CD28 antibodies.
(**I**) Changes in Ki-$67^+$ cell frequency after ICI therapy in DP-Tem, SP-Tem, and DN-Tem (n=20).
All bar graphs represent mean ± SD, ∗∗∗$p<0.001$, ∗∗$p<0.01$, ∗$p<0.05$.

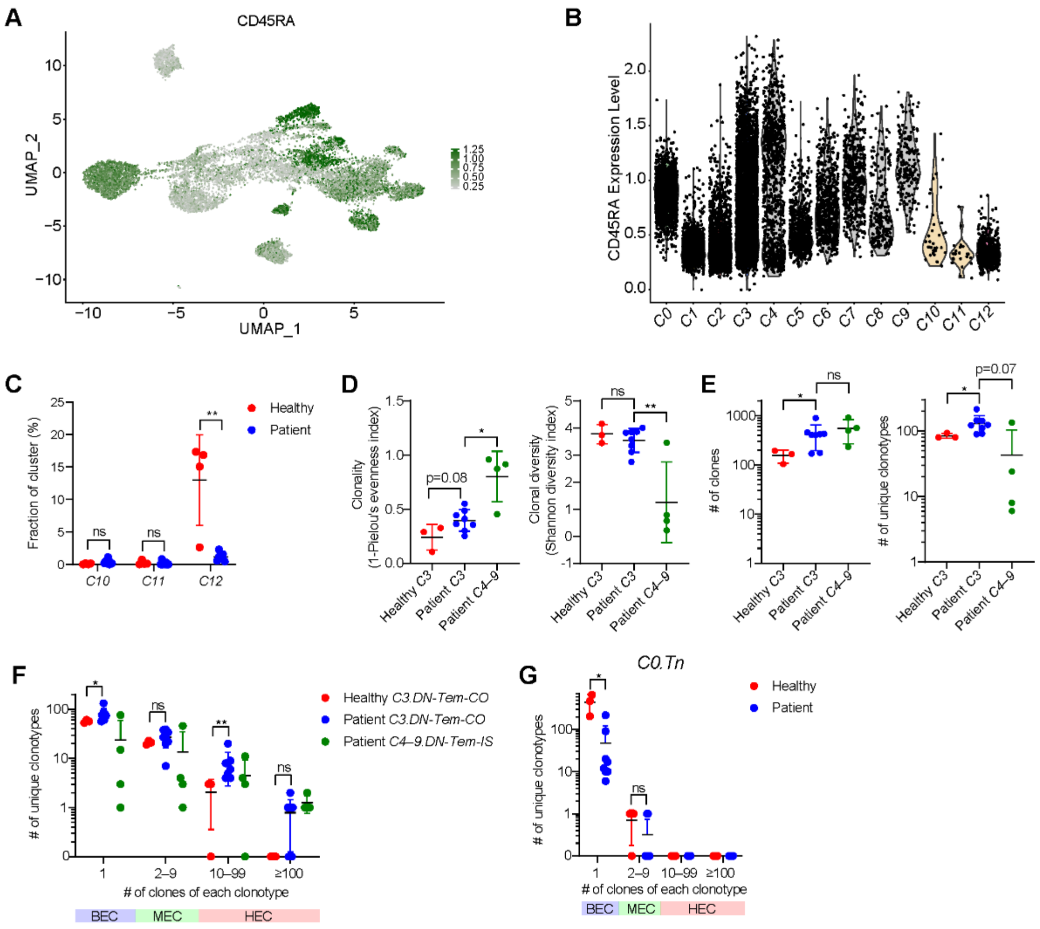


**Fig. S3. *DN-Tem* clusters in NSCLC patients exhibit an increased number of unique clonotypes.**

(**A**) UMAP of CD45RA expression measured using CITE-seq.

(**B**) CD45RA expression in each cluster.

(**C**) Frequencies of *C10–12* clusters in healthy individuals (n=4) and patients (n=8).

(**D**–**E**) (D) Clonality, clonal diversity, and (E) numbers of clones and unique clonotypes in *C3* and *C4–9* clusters of healthy individuals (n=3) and patients (n=8).

(**F**–**G**) Number of unique clonotypes in (F) *C3.DP-Tem-CO*, *C4–9.DN-Tem-IS*, and (G) *C0.Tn* clusters that have only one clone (barely expanded clones; BEC), 2–9 clones (moderately expanded clones; MEC), 10–99 or ≥100 clones (heavily expanded clones; HEC).

All bar graphs represent mean ± SD, ∗∗p<0.01, ∗p<0.05.

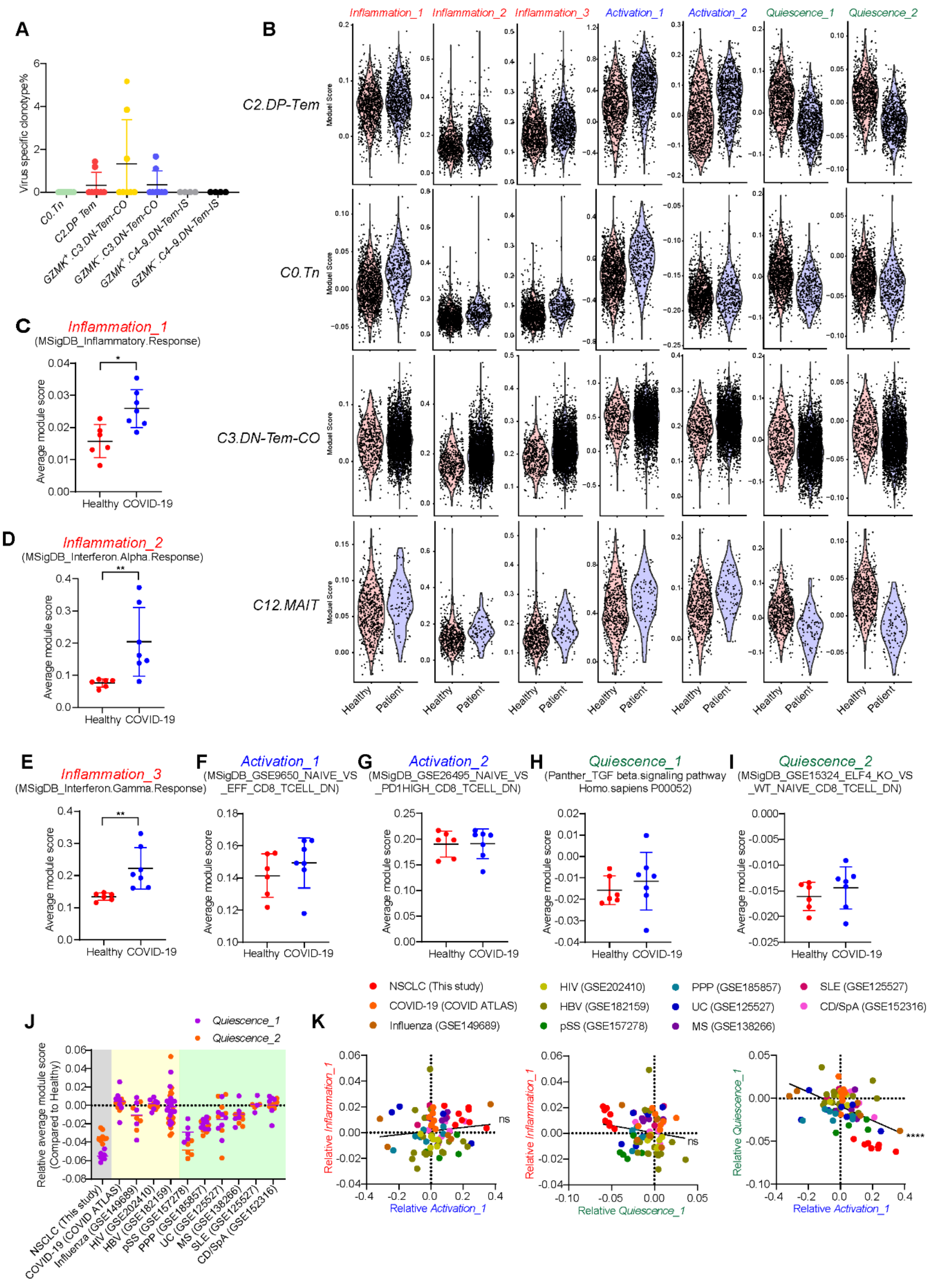


**Fig. S4. T cell quiescence signatures inversely correlate with T cell activation signatures in peripheral blood CD8$^{+}$ T cells of NSCLC patients.**

(**A**) Frequency of reported virus-specific unique clonotypes in each cluster.
(**B**) Module scores for each gene set in *C2.DP-Tem, C0.Tn, C3-DN-Tem-CO,* and *C12.MAIT* clusters.
(**C–I**) Average module scores of indicated gene sets in *DP-Tem* (*GZMK*$^{+}$*GZMH*$^{-}$) cluster of healthy individuals (n=6) and COVID-19 patients (n=7).
(**J**) Relative average module score of *Quiescence_1* and *Quiescence_2* gene sets in patients with NSCLC (n=8), COVID-19 (n=7), influenza (n=5), human immunodeficiency virus (HIV; n=6), hepatitis B virus (HBV; n=18), progressive Sjögren's syndrome (pSS; n=5), palmoplantar pustulosis (PPP; n=7), ulcerative colitis (UC; n=7), multiple sclerosis (MS; n=5), systemic lupus erythematosus (SLE; n=3), Crohn's disease, or spondyloarthritis (CD and SpA, respectively; n=6). Relative average module score was calculated by subtracting average module scores of patients with that of healthy controls from the same dataset in *DP-Tem* (*GZMK*$^{+}$*GZMH*$^{-}$*)* clusters.
(**K**) Correlation between signature gene sets. Dotted lines represent the average module scores of healthy controls. Solid lines represent linear regression, with p-values calculated using an F-test for non-zero correlation.
All bar graphs represent mean ± SD. ∗∗∗∗p<0.0001, ∗∗p<0.01, ∗p<0.05.

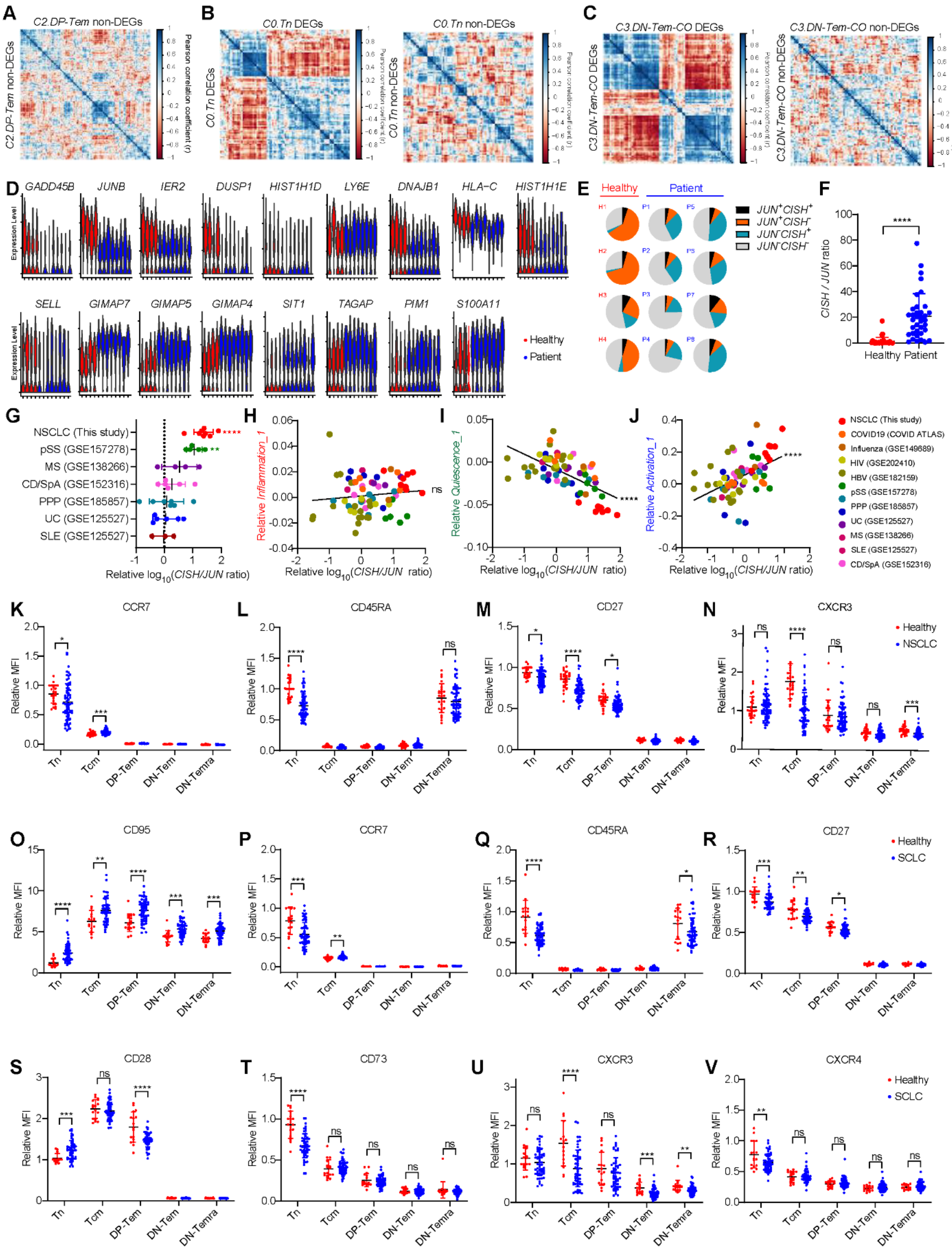


**Fig. S5. Peripheral blood CD8$^{+}$ T cells in NSCLC patients exhibit coordinated gene and protein expression alterations indicative of systemic homeostatic dysregulation.**

(**A**) Correlation matrix of 150 randomly selected non-DEGs of *C2.DP-Tem*. Pearson correlation coefficient was color coded from red (low) to blue (high).
(**B**) Correlation matrix of 92 *C0.Tn* DEGs (left) and 92 randomly selected *C0.Tn* non-DEGs (right).
(**C**) Correlation matrix of 84 *C3.DN-Tem-CO* DEGs (left) and 84 randomly selected *C3.DN-Tem-CO* non-DEGs (right).
(**D**) Expressions of DEGs that were commonly observed (except *CISH* and *JUN*) in the three clusters (*C0.Tn, C2.DP-Tem,* and *C3.DN-Tem-CO*).
(**E**) Pie chart representing *JUN* and/or *CISH* expressing cells in total $CD8^+$ T cells in each individual.
(**F**) *CISH/JUN* ratio by quantitative PCR using whole peripheral blood mononuclear cells of healthy individuals (n=37) and NSCLC patients (n=36).
(**G**) Relative $\log_{10}$(*CISH/JUN* ratio) of patients with NSCLC or autoimmune diseases. Relative $\log_{10}$(*CISH/JUN* ratio) was calculated by subtracting the $\log_{10}$(*CISH/JUN* ratio) of patients with the average $\log_{10}$(*CISH/JUN* ratio) of healthy individuals within *DP-Tem* (*$GZMK^+GZMH^-$*) clusters. The dotted line represents the average $\log_{10}$(*CISH/JUN* ratio) of healthy individuals.
(**H**–**J**) Correlations between $\log_{10}$(*CISH/JUN* ratio) and signature gene sets. Lines represent linear regression, with p-values calculated using an F-test for non-zero correlation
(**K**–**N**) Relative MFIs of indicated molecules in each subset from healthy individuals (n=25) and NSCLC patients (n=71). Relative MFIs were calculated by dividing MFIs with MFI of Tn from a healthy individual.
(**O**–**V**) Relative MFIs of indicated molecules in each subset from healthy individuals (n=17) and SCLC patients (n=55).
All bar graphs represent mean ± SD. ∗∗∗∗$p<0.0001$, ∗∗∗$p<0.001$, ∗∗$p<0.01$, ∗$p<0.05$.

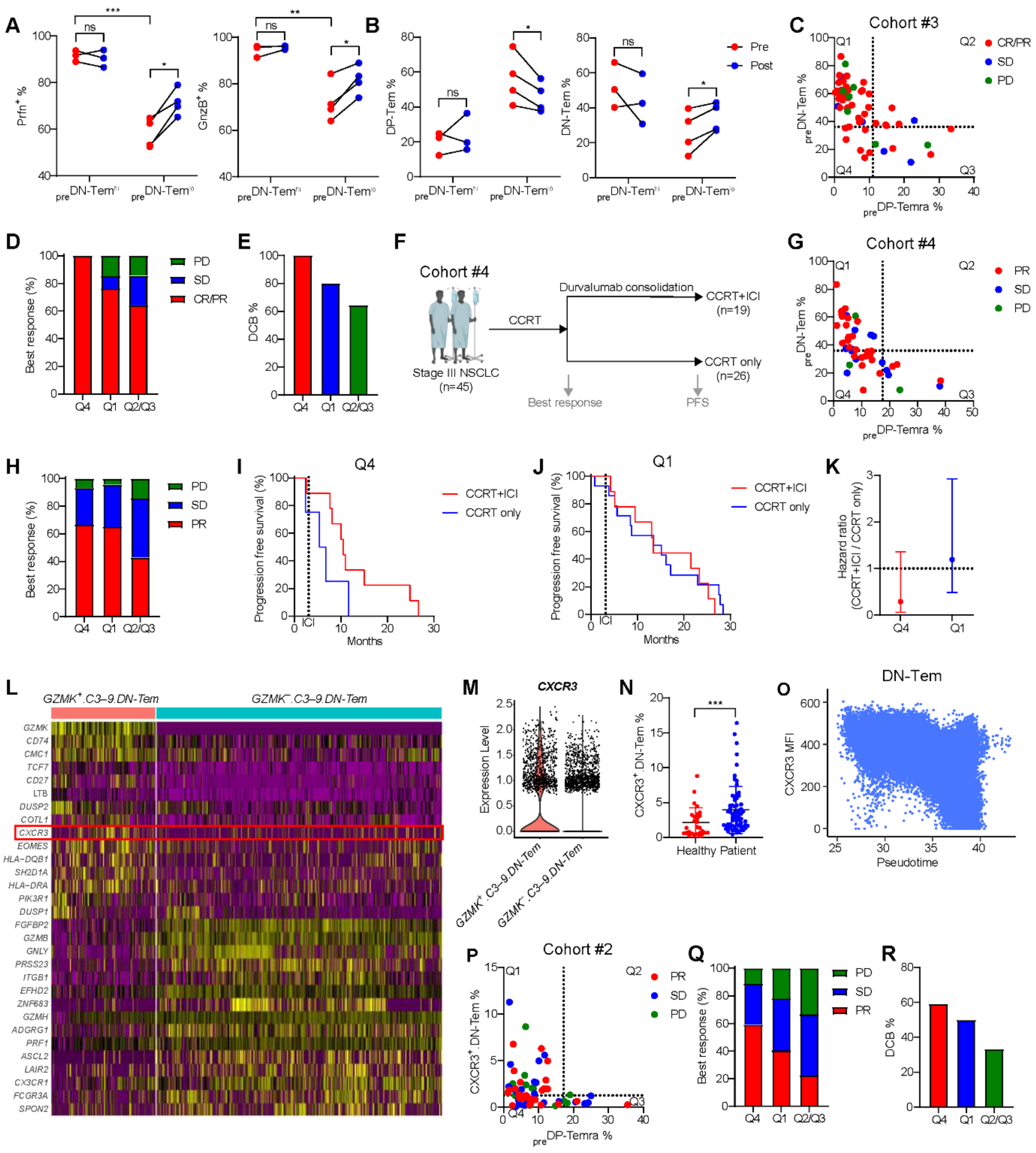

**Fig. S6. Two subsets of peripheral blood CD8$^+$ T cells predict clinical outcomes to ICI therapy.**

(**A**–**B**) Changes in (A) prfn$^+$ and gnzB$^+$ frequencies and (B) DP-Tem and DN-Tem frequencies after pembrolizumab treatment in $_{pre}$DN-Tem$^{hi}$ (n=3) and $_{pre}$DN-Tem$^{lo}$ groups (n=4). The bar graphs represent mean ± SD.

(**C**–**E**) Relationship between frequencies of $_{pre}$DN-Tem and $_{pre}$DP-Temra and (C–D) complete response (CR)/PR and (E) durable clinical benefit (DCB) in Cohort #3 (n=55).

(**F**) Treatment strategy for Cohort #4.

(**G**–**H**) Relationship between frequencies of $_{pre}$DN-Tem and $_{pre}$DP-Temra and best response to concurrent chemo-radiation therapy (CCRT) in Cohort #4 (n=45).
(**I**–**J**) Progression free survival (PFS) of patients treated with CCRT alone (CCRT only) or CCRT and durvalumab consolidation (CCRT+ICI) among patients in (I) Q4 or (J) Q1.
(**K**) Hazard ratio for PFS in CCRT+ICI patients, with reference to CCRT only patients, among the patients in Q4 or Q1. The error bars represent 95% confidence interval.
(**L**) Heat map for DEGs between *GZMK*$^{+}$*.C3–9.DN-Tem* and *GZMK*$^{-}$*.C3–9.DN-Tem* in scRNA-seq data. Expressions were color coded from purple (low) to yellow (high).
(**M**) Gene expression of *CXCR3* in *GZMK*$^{+}$*.C3–9.DN-Tem* and *GZMK*$^{-}$*.C3–9.DN-Tem.*
(**N**) Frequency of CXCR3$^{+}$.DN-Tem in CD8$^{+}$ T cells from healthy individuals (n=28) or stage IV NSCLC patients (n=68) analyzed by flow cytometry. The bar graph represents mean ± SD.
(**O**) CXCR3 expression kinetics in DN-Tem over pseudotimes generated with protein expressions.
(**P**–**R**) Relationship between frequencies of CXCR3$^{+}$.DN-Tem and $_{pre}$DP-Temra and (P-Q) PR and (R) DCB in Cohort #2 (n=68).
∗∗∗p<0.001, ∗∗p<0.01, ∗p<0.05.

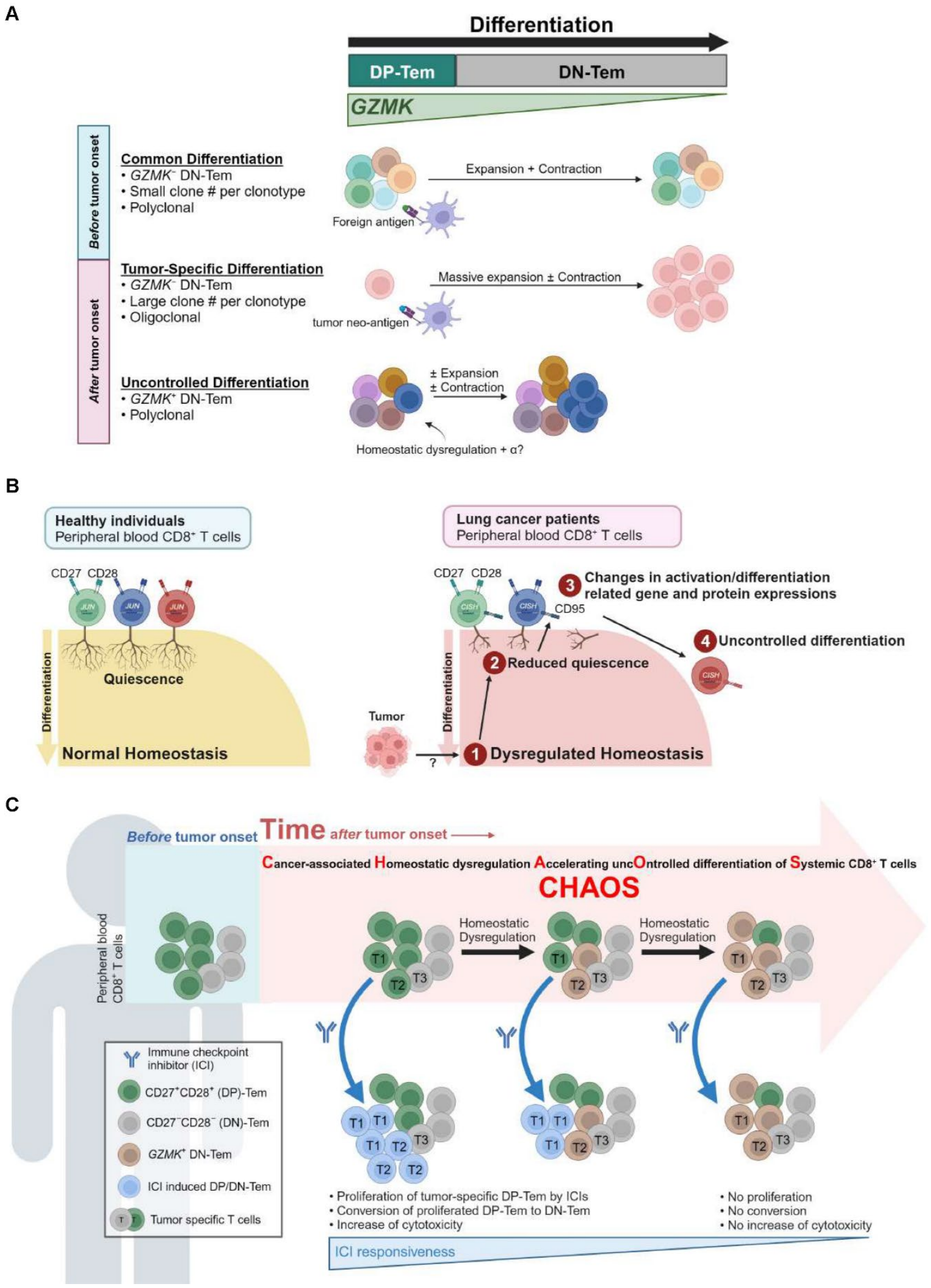


**Fig. S7. Cancer-associated homeostatic dysregulation accelerates uncontrolled differentiation of systemic CD8⁺ T cells.**

(**A**) Proposed model for heterogenous differentiation pathways in generating DN-Tem. The common differentiation pathway is shared by both healthy individuals and cancer patients, producing *GZMK⁻.DN-Tem* that are polyclonal and typically have a small number of clones. The tumor-specific differentiation pathway is exclusive to cancer patients, generating *GZMK⁻.DN-Tem* that are oligoclonal but have a high number of clones, suggesting they may be generated in response to tumor neo-antigens. The uncontrolled differentiation pathway, also found exclusively in cancer patients, generates *GZMK⁺.DN-Tem* that are highly diverse (polyclonal) and exhibit varying numbers of clones. These cells are induced by homeostatic dysregulation.
(**B**) Proposed model for uncontrolled differentiation of systemic $CD8^+$ T cells. In healthy individuals, T cell homeostasis is well-maintained, where T cell quiescence prevents aberrant activation of $CD8^+$ T cells. However, when tumors develop, they induce a systematic and systemic dysregulation in T cell homeostasis through yet unknown mechanisms. This dysregulation leads to decreased T cell quiescence, which in turn alters various gene and protein expressions putatively involved in the regulation of T cell activation and differentiation, including *JUN* and *CISH*. These changes ultimately lead to a stepwise, uncontrolled transition to a more-differentiated state in all peripheral blood $CD8^+$ T cell populations.
(**C**) Proposed model "Cancer-associated Homeostatic dysregulation Accelerating uncOntrolled differentiation of Systemic $CD8^+$ T cells" (CHAOS). In the earlier times after tumor onset, tumor-specific DP-Tem respond to ICI therapy, leading to their proliferation and differentiation. This process results in an increased frequency of DN-Tem and enhanced cytotoxic activity, resulting in a favorable response to ICI therapy. However, in the later times after tumor onset, CHAOS induces a progressive transition of systemic $CD8^+$ T cells from DP-Tem to DN-Tem (particularly *GZMK⁺.DN-Tem*). Tumor-specific $CD8^+$ T cells that underwent DP-to-DN-Tem transition by CHAOS exhibit impaired responsiveness to ICI therapy, ultimately leading to poor clinical outcomes.

| Characteristics | | Number (n=349) | Proportion (%) |
|---|---|---|---|
| **Age** | 30s | 2 | 0.6 |
| | 40s | 13 | 3.7 |
| | 50s | 34 | 9.7 |
| | 60s | 137 | 39.3 |
| | 70s | 136 | 39.0 |
| | 80s | 23 | 6.6 |
| | NA | 4 | 1.1 |
| **Sex** | Male | 288 | 82.5 |
| | Female | 58 | 16.6 |
| | NA | 3 | 0.9 |
| **Smoking** | Smoker | 265 | 75.9 |
| | Never-smoker | 62 | 17.8 |
| | NA | 22 | 6.3 |
| **Stage** | Stage I NSCLC | 33 | 9.5 |
| | Stage II NSCLC | 4 | 1.1 |
| | Stage III NSCLC | 60 | 17.2 |
| | Stage IV NSCLC | 197 | 56.4 |
| | Stage ED SCLC | 55 | 15.8 |
| **Tumor PD-L1** | 0% | 136 | 39.0 |
| | 1–49% | 60 | 17.2 |
| | 50–100% | 68 | 19.5 |
| | NA | 85 | 24.4 |
| **Histology** | SQC NSCLC | 111 | 31.8 |
| | Non-SQC NSCLC | 179 | 51.3 |
| | SCLC | 55 | 15.8 |
| | NA | 4 | 1.1 |
| **EGFR mutation** | Wildtype | 189 | 54.2 |
| | E19del | 12 | 3.4 |
| | L858R | 13 | 3.7 |
| | S768I | 1 | 0.3 |
| | E19del/T790M | 2 | 0.6 |
| | L858R/T790M | 1 | 0.3 |
| | NA | 131 | 37.5 |

**Table S1. Patient information**

NA, not available; NSCLC, non-small cell lung cancer; SCLC, small cell lung cancer; LD, limited disease; ED, extensive disease; PD-L1, programmed cell death ligand-1; SQC, squamous cell carcinoma; EGFR, epidermal growth factor.

| Characteristics | | Number (n=32) | Proportion (%) |
|---|---|---|---|
| **Age** | 50s | 7 | 21.9 |
| | 60s | 17 | 53.1 |
| | 70s | 5 | 15.6 |
| | 80s | 3 | 9.4 |
| **Sex** | Male | 29 | 90.6 |
| | Female | 3 | 9.4 |
| **Smoking** | Smoker | 28 | 87.5 |
| | Never-smoker | 4 | 12.5 |
| **Stage** | Stage IV NSCLC | 32 | 100.0 |
| **Tumor PD-L1** | 0% | 14 | 43.8 |
| | 1–49% | 3 | 9.4 |
| | 50–100% | 12 | 37.5 |
| | NA | 3 | 9.4 |
| **Histology** | SQC NSCLC | 13 | 40.6 |
| | Non-SQC NSCLC | 19 | 59.4 |
| **EGFR mutation** | Wildtype | 18 | 56.3 |
| | E19del | 3 | 9.4 |
| | L858R | 1 | 3.1 |
| | NA | 10 | 31.3 |
| **ICI** | Atezolizumab | 32 | 100.0 |
| **IO tx line** | 2$^{nd}$ | 21 | 65.6 |
| | 3$^{rd}$ | 6 | 18.8 |
| | >4$^{th}$ | 5 | 15.6 |
| **Post-therapy collection date** | 7 | 12 | 37.5 |
| | 8 | 3 | 9.4 |
| | 9 | 2 | 6.3 |
| | 10 | 10 | 31.3 |
| | 11 | 1 | 3.1 |
| | 14 | 4 | 12.5 |
| **Best response** | PR | 6 | 18.8 |
| | SD | 9 | 28.1 |
| | PD | 16 | 50.0 |
| | NE | 1 | 3.1 |

**Table S2. Patient information**
NA, not available; NSCLC, non-small cell lung cancer; PD-L1, programmed cell death ligand-1; SQC, squamous cell carcinoma; EGFR, epidermal growth factor; ICI, immune checkpoint inhibitor; IO, immuno-oncology; tx, therapy; PR, partial response; SD, stable disease; PD, progressive disease; NE, inevaluable.

| Patient | P1 | P2 | P3 | P4 | P5 | P6 | P7 | P8 |
|---|---|---|---|---|---|---|---|---|
| **Age** | 72 | 64 | 66 | 58 | 67 | 62 | 44 | 66 |
| **Sex** | M | M | M | F | M | M | M | M |
| **Smoking** | Smoker | Smoker | Smoker | Never | Smoker | Smoker | Smoker | Smoker |
| **Stage** | IB | IA3 | IB | IA2 | IVB | IVB | IVB | IVA |
| **Histology** | ADC | ADC | SQC | ADC | ADC | ADC | ADC | ADC |
| **EGFR mutation** | L858R | WT | WT | L858R | WT | WT | WT | L858R |
| **Tumor PD-L1** | 90 | 100 | 5 | 0 | 50 | 80 | 15 | 60 |

**Table S3. Patient information for scRNA-seq**
ADC, adenocarcinoma; SQC, squamous cell carcinoma; EGFR, epidermal growth factor; WT, wildtype; PD-L1, programmed cell death ligand-1.

| Characteristics | | Number (n=92) | Proportion (%) |
|---|---|---|---|
| **Age** | 40s | 5 | 5.4 |
| | 50s | 17 | 18.5 |
| | 60s | 43 | 46.7 |
| | 70s | 22 | 23.9 |
| | 80s | 5 | 5.4 |
| **Sex** | Male | 74 | 80.4 |
| | Female | 18 | 19.6 |
| **Smoking** | Smoker | 72 | 78.3 |
| | Never-smoker | 20 | 21.7 |
| **Stage** | Stage IV NSCLC | 92 | 100.0 |
| **Tumor PD-L1** | 0% | 50 | 54.3 |
| | 1–49% | 18 | 19.6 |
| | 50–100% | 19 | 20.7 |
| | NA | 5 | 5.4 |
| **Histology** | SQC NSCLC | 32 | 34.8 |
| | Non-SQC NSCLC | 60 | 65.2 |
| **EGFR mutation** | Wildtype | 55 | 59.8 |
| | E19del | 9 | 9.8 |
| | L858R | 4 | 4.3 |
| | S768I | 1 | 1.1 |
| | L858R/T790M | 1 | 1.1 |
| | NA | 22 | 23.9 |
| **Therapy** | Atezolizumab | 92 | 100.0 |
| **IO tx line** | 2$^{nd}$ | 54 | 58.7 |
| | 3$^{rd}$ | 16 | 17.4 |
| | >4$^{th}$ | 22 | 23.9 |
| **Best response** | PR | 14 | 15.2 |
| | SD | 28 | 30.4 |
| | PD | 50 | 54.3 |
| **Durable benefit** | DCB | 22 | 23.9 |
| | NCB | 67 | 72.8 |
| | NA | 3 | 3.3 |
| **PFS** | 0–5 months | 67 | 72.8 |
| | 5–10 months | 12 | 13.0 |
| | 10–15 months | 4 | 4.3 |
| | 15–20 months | 1 | 1.1 |
| | 20–25 months | 5 | 5.4 |
| | NA | 3 | 3.3 |

**Table S4. Patient information for Cohort #1**
NA, not available; NSCLC, non-small cell lung cancer; PD-L1, programmed cell death ligand-1; SQC, squamous cell carcinoma; EGFR, epidermal growth factor; IO, immuno-oncology; tx, therapy; PR, partial response; SD, stable disease; PD, progressive disease; DCB, durable clinical benefit; NCB, no durable clinical benefit; PFS, progression free survival.

| **Characteristics** | | **Number (n=87)** | **Proportion (%)** |
|---|---|---|---|
| **Age** | 30s | 1 | 1.1 |
| | 40s | 4 | 4.6 |
| | 50s | 6 | 6.9 |
| | 60s | 30 | 34.5 |
| | 70s | 39 | 44.8 |
| | 80s | 7 | 8.0 |
| **Sex** | Male | 77 | 88.5 |
| | Female | 10 | 11.5 |
| **Smoking** | Smoker | 72 | 82.8 |
| | Never-smoker | 15 | 17.2 |
| **Stage** | Stage IV NSCLC | 87 | 100.0 |
| **Tumor PD-L1** | 0% | 35 | 40.2 |
| | 1–49% | 18 | 20.7 |
| | 50–100% | 32 | 36.8 |
| | NA | 2 | 2.3 |
| **Histology** | SQC NSCLC | 29 | 33.3 |
| | Non-SQC NSCLC | 58 | 66.6 |
| **EGFR mutation** | WT | 62 | 71.3 |
| | L858R | 2 | 2.3 |
| | E19del/T790M | 1 | 1.1 |
| | NA | 22 | 25.3 |
| **Therapy** | K | 26 | 29.9 |
| | KAP | 28 | 32.2 |
| | KAC | 13 | 14.9 |
| | KTC | 20 | 23.0 |
| **IO tx line** | $1^{st}$ | 69 | 79.3 |
| | $2^{nd}$ | 11 | 12.6 |
| | $3^{rd}$ | 5 | 5.7 |
| | $4^{th}$ | 2 | 2.3 |
| **Best response** | PR | 34 | 39.1 |
| | SD | 27 | 31.0 |
| | PD | 26 | 29.9 |
| **Durable benefit** | DCB | 40 | 46.0 |
| | NCB | 47 | 54.0 |

**Table S5. Patient information for Cohort #2**
NA, not available; NSCLC, non-small cell lung cancer; PD-L1, programmed cell death ligand-1; SQC, squamous cell carcinoma; EGFR, epidermal growth factor; K, pembrolizumab alone; KAP, pembrolizumab, pemetrexed, and cisplatin combination therapy; KAC, pembrolizumab, pemetrexed, and carboplatin combination therapy; KTC, pembrolizumab, paclitaxel, and carboplatin combination therapy; IO, immuno-oncology; tx, therapy; PR, partial response; SD, stable disease; PD, progressive disease; DCB, durable clinical benefit; NCB, no durable clinical benefit.

| **Characteristics** | | **Number (n=55)** | **Proportion (%)** |
|---|---|---|---|
| **Age** | 50s | 1 | 1.8 |
| | 60s | 19 | 34.5 |
| | 70s | 25 | 45.5 |
| | 80s | 10 | 18.2 |
| **Sex** | Male | 52 | 94.5 |
| | Female | 3 | 5.5 |
| **Smoking** | Smoker | 48 | 87.3 |
| | Never-smoker | 7 | 12.7 |
| **Stage** | Stage ED SCLC | 55 | 100.0 |
| **Histology** | SCLC | 55 | 100.0 |
| **Therapy** | ACE | 55 | 100.0 |
| **IO tx line** | 1st | 55 | 100.0 |
| **Best response** | CR | 1 | 1.8 |
| | PR | 41 | 74.5 |
| | SD | 6 | 10.9 |
| | PD | 5 | 9.1 |
| | NA | 2 | 3.6 |
| **Durable benefit** | DCB | 43 | 78.2 |
| | NCB | 12 | 21.8 |

**Table S6. Patient information for Cohort #3**
NA, not available; ED, extensive disease; SCLC, small cell lung cancer; ACE, atezolizumab, carboplatin, and etoposide combination therapy; IO, immuno-oncology; tx, therapy; CR, complete response; PR, partial response; SD, stable disease; PD, progressive disease; DCB, durable clinical benefit; NCB, no durable clinical benefit.

| Characteristics | | Number (n=45) | Proportion (%) |
|---|---|---|---|
| **Age** | 50s | 7 | 15.6 |
| | 60s | 16 | 35.6 |
| | 70s | 21 | 46.7 |
| | 80s | 1 | 2.2 |
| **Sex** | Male | 40 | 88.9 |
| | Female | 5 | 11.1 |
| **Smoking** | Smoker | 41 | 91.1 |
| | Never-smoker | 4 | 8.9 |
| **Stage** | Stage III NSCLC | 45 | 100.0 |
| **Tumor PD-L1** | 0% | 18 | 40.0 |
| | 1–49% | 15 | 33.3 |
| | 50–100% | 12 | 26.7 |
| **Histology** | SQC NSCLC | 26 | 57.8 |
| | Non-SQC NSCLC | 19 | 42.2 |
| **EGFR mutation** | WT | 19 | 42.2 |
| | L858R | 2 | 4.4 |
| | NA | 24 | 53.3 |
| **1st line therapy** | CCRT | 45 | 100.0 |
| **Best response** | PR | 28 | 62.2 |
| | SD | 14 | 31.1 |
| | PD | 3 | 6.7 |
| **Consolidation** | Durvalumab | 19 | 42.2 |
| | None | 26 | 57.8 |
| **PFS** | 0–5 months | 6 | 13.3 |
| | 5–10 months | 12 | 26.7 |
| | 10–15 months | 7 | 15.6 |
| | 15–20 months | 5 | 11.1 |
| | 20–25 months | 6 | 13.3 |
| | 25–30 months | 8 | 17.8 |
| | NA | 1 | 2.2 |

**Table S7. Patient information for Cohort #4**
NA, not available; NSCLC, non-small cell lung cancer; PD-L1, programmed cell death ligand-1; SQC, squamous cell carcinoma; EGFR, epidermal growth factor; CRT, concurrent chemoradiation therapy; PR, partial response; SD, stable disease; PD, progressive disease; PFS, progression free survival.
